# Serum Lipidome Profiling Reveals a Distinct Signature of Ovarian Cancer in Korean Women

**DOI:** 10.1101/2023.10.05.560751

**Authors:** Samyukta Sah, Olatomiwa O. Bifarin, Samuel G. Moore, David A. Gaul, Hyewon Chung, Hanbyoul Cho, Chi-Heum Cho, Jae-Hoon Kim, Jaeyeon Kim, Facundo M. Fernández

## Abstract

Distinguishing ovarian cancer (OC) from other gynecological malignancies remains a critical unmet medical need with significant implications for patient survival. However, non-specific symptoms along with our lack of understanding of OC pathogenesis hinder its diagnosis, preventing many women from receiving appropriate medical assistance. Accumulating evidence suggests a link between OC and deregulated lipid metabolism. Most studies, however, are limited by small sample size, particularly for early-stage cases. Furthermore, racial/ethnic differences in OC survival and incidence have been reported, yet most of the studies consist largely of non-Hispanic white women or women with European ancestry. Studies of more diverse racial/ethnic populations are needed to make OC diagnosis and prevention more inclusive. Here, we profiled the serum lipidome of 208 OC, including 93 patients with early-stage OC, and 117 non-OC (other gynecological malignancies) patients of Korean descent. Serum samples were analyzed with a high-coverage liquid chromatography high-resolution mass spectrometry platform, and lipidome alterations were investigated *via* statistical and machine learning approaches. Results show that lipidome alterations unique to OC were present in Korean women as early as when the cancer is localized, and those changes increase in magnitude as the diseases progresses. Analysis of relative lipid abundances revealed specific patterns for various lipid classes, with most classes showing decreased abundance in OC in comparison to other gynecological diseases. Machine learning methods selected a panel of 17 lipids that discriminated OC from non-OC cases with an AUC of 0.85 for an independent test set. This study provides a systemic analysis of lipidome alterations in human OC, specifically in Korean women, emphasizing the potential of circulating lipids in distinguishing OC from non-OC conditions.

## Introduction

The high mortality rate of ovarian cancer (OC) is largely due to the asymptomatic progression of the disease, accompanied by our lack of understanding of OC biology. OC is not a single disease; it consists of several histological subtypes^1^ and its heterogenous nature is one major obstacle in understanding disease biology and identifying new biomarkers^2^. Presently, there are no screening tests available for detecting OC in the general population^3^. In many cases, women with OC experience non-specific symptoms such as pelvic pain and bloating, and thus are frequently dismissed and misdiagnosed as benign conditions^4^. Symptomatic patients with a risk of developing OC are screened via trans-vaginal sonography (TVS) or the measurement of serum protein biomarker CA125^5^. However, these diagnostic strategies lack adequate sensitivity and specificity. For instance, increased levels of CA125 are observed for several non-OC related gynecological malignancies while it is only elevated in 50% of early OC cases^5^. Likewise, TVS can misdiagnose OC as benign diseases, leading to mismanagement that significantly reduces patient survival rate^6^. OC patients misdiagnosed for benign diseases suffer worse prognosis compared to women who are treated by gynecologic oncologists^7^, and currently 30-50% of women with OC in the United States do not receive appropriate OC treatment^4^. Thus, while effective screening in the general population remains the goal, accurate diagnosis and triage of women suspected of OC is crucial for improved prognosis.

Substantial effort has been put into improving the clinical diagnosis of OC and understanding the underlying disease mechanisms. For example, the two-stage approach that uses both CA125 and TVS sequentially showed improved specificity in clinical trials, and the use of protein biomarker HE4 in combination with CA125 has shown better specificity for distinguishing malignant from benign pelvic masses^8^. Other potential biomarkers for improved diagnosis include autoantibodies and antigen-autoantibody complexes, sometimes combined with CA125^9, 10^, serum microRNAs^11^or circulating tumor DNA^12^. Recently, protein markers detected in Pap test fluid^13^ have shown promising results to distinguish OC from women with normal cytology. One characteristic feature of malignant tumors is their ability to rewire metabolism in response to high cell proliferation rates^14^. Metabolomics, the examination of metabolites including lipids, carbohydrates, and amino acids, provides a powerful platform for investigating metabolic alterations and accelerating biomarker discovery. As metabolomics studies generate hundreds of thousands of spectral signals, machine learning (ML) is indispensable to metabolomics data analysis^15^.

In recent years, alterations in ovarian cancer lipid metabolism have gained increased attention^16, 17^. Lipids function as building blocks for cell membranes, participate in cellular signaling, and are regulators of numerous cellular functions that drive energy-related processes^18^. Given the close connection between altered lipid metabolism and oncogenesis, there is accumulating evidence showing specific lipid profiles associated with OC growth and metastasis^16, 19, 20^. Serum lipidome profiling of ovarian cancer patients against normal controls and benign malignancies has shown evidence of dysregulation in glycerophospholipids, ceramides, and triglycerides being associated with malignant ovarian tumors^16, 17, 21^. Some studies have suggested the use of lipid panels in combination with CA125 can achieve enhanced diagnostic power^17, 22^. The diagnostic potential of gangliosides, a class of glycolipids involved in immunosuppressive response in tumors, has recently been reported for distinguishing OC patients from non-OC related diseases as well as healthy controls^23^. Increased ganglioside levels in plasma, tissue, and ascites fluid from OC patients have also been reported^23, 24^. These studies, however, are often hindered by small sample sizes, limited number of early-stage samples, lack of external validation datasets, and inconsistencies in sample collection and processing protocols, thereby limiting the statistical significance and robustness of the final results. Most studies focus on cohorts of non-Hispanic white women or women with European ancestry, and a few examples involving cohorts of Chinese women^25, 26^. However, studies with patients of Korean descent are largely lacking. Although an integrated serum proteomics and metabolomics study of Korean patients has been previously reported, it only involved 10 OC patients^27^.

Here, we present a comprehensive serum lipidomic study of ovarian cancer patients of Korean descent with various histological types and disease stages (*n* = 208) and of women with other gynecological malignancies, including invasive cervical cancer (*n* =117). Samples were secured from two independent tissue banks. Ultrahigh performance liquid chromatography-high resolution mass spectrometry (UHPLC-HRMS) combined with machine learning (ML) were used to identify lipidome alterations that distinguished between OC and non-OC patients. One thousand and sixteen lipid species belonging to 23 lipid sub classes were detected and compared between OC and non-OC cases, providing a systems level perspective of OC-related lipidome perturbations. Stage-stratified analysis revealed lipid alterations specific to early or advanced stages in comparison to non-OC conditions. An ML based pipeline was developed and a panel of 10 best lipids was selected. This panel discriminated OC *vs.* non-OC cases with area under curve (AUC) of 0.85 and 0.82 for the cross-validated training and test sets, respectively. Considering the emerging potential of gangliosides as circulating markers for OC, a set of optimal discriminating gangliosides was also selected and incorporated into the 10-lipid biomarker panel, achieving slightly improved AUC values of 0.88 and 0.83 for training and test sets, respectively. In addition to traditional ML approaches, we applied an AutoML method, Auto-sklearn^28^, to further assess the classification performance of the 17-lipid panel. Improved classification metrics were achieved with the AutoML method with an AUC of 0.85 for an independent test set.

## Experimental Section

### Patient Cohort

Serum samples were obtained from two independent tissue banks in South Korea: Dongsan Hospital Human Tissue Bank and the Human Tissue Bank of Gangnam Severance Hospital, Yonsei University College of Medicine (No. HTB-P2019-13). Samples from both tissue banks were obtained after the approval from their respective IRB and the patient’s informed consent. The Severance cohort included 185 samples from ovarian cancer patients, 47 from women with benign ovarian tumors, 50 from invasive cervical cancers and 21 samples from patients with benign uterine tumors. Blood was collected from all patients during surgery after anesthesia and at least 8 hours of fasting. In the Dongsan cohort, 88 women had ovarian cancer, 12 had benign ovarian tumors, 10 had benign uterine tumors, and 9 women had cervical cancer. As with the Severance cohort, samples from these patients were collected during surgery after anesthesia and at least 6 hours of fasting. All recruited participants were of Korean descent. Samples from both cohorts were grouped together and patients with ovarian cancer (OC) and all other gynecological malignancies (non-OC) were age matched. The matched cohort included 208 patients with ovarian cancer (mean age 51.9 years) and 117 non-OC patients (mean age 49.9 years). Disease stages and histological characteristics of each patient are given in **Table 1**. Among the OC patients, 93 patients had early stage (I and II) cancers. Ten of the OC patients had recurrent cancer and 9 have had their samples collected more than once. Samples from normal controls (*i.e*., women with no known gynecological malignancies) were also collected during regular health exams at Severance hospital. In this case, blood was collected after fasting for at least 8 hours. As noted earlier, blood from OC patients or patients with other conditions was collected after the initiation of anesthesia, and thus could not be directly compared with normal controls without major confounding effects. Therefore, control samples were excluded from the main data analysis pipeline. However, we also conducted a lipidome comparison study between healthy controls and OC patients for reference purposes. Results from this study are presented in the supplemental information section and the data are shared together with the rest of the cohort.

**Table 1.**
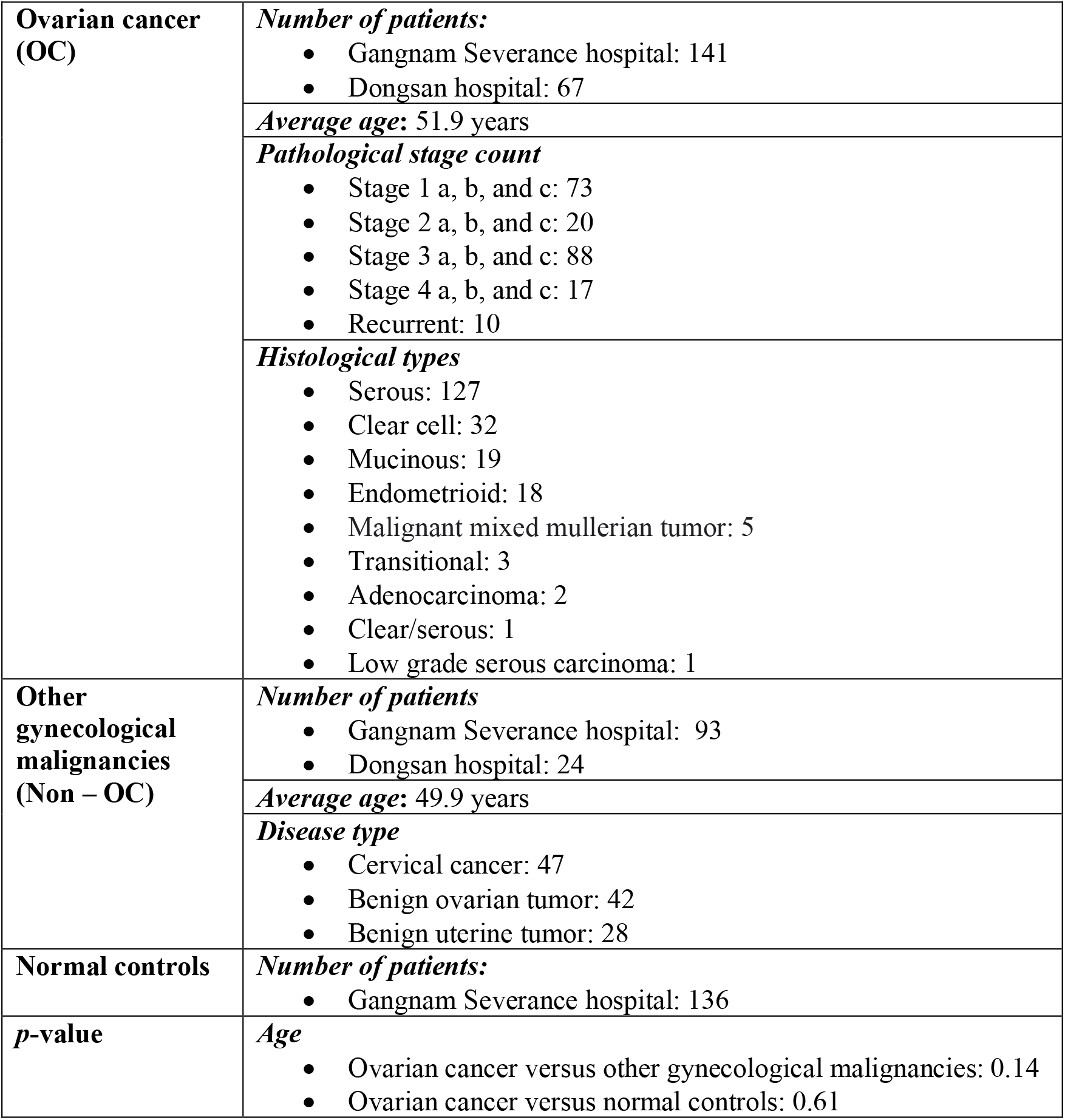
Matched Cohort Characteristics. Patients from OC and non-OC groups were matched by age. The matched cohort included *n* = 208 OC and *n* = 117 non-OC serum samples. All *p*-values were calculated using Welch’s t-test.

### Chemicals

LC-MS grade 2-propanol, water, formic acid (99.5+ %), ammonium formate and ammonium acetate were purchased from Fisher Chemical (Fisher Scientific International, Inc. Pittsburgh, PA) and used for the preparation of chromatographic mobile phases and sample extraction. Isotopically labeled lipid standards (**Table S1**) were purchased from Avanti Polar Lipids (Alabaster, AL, USA).

### Sample Preparation

Serum samples were thawed on ice, followed by extraction of the non-polar (lipid) metabolome. The extraction solvent was prepared by addition of 725 *μ*L of the isotopically labeled lipid standard mixture (**Table S1**) to 43.5 mL 2-propanol (1:60 ratio) and kept on ice. This cold extraction mixture was added to serum samples in a solvent: serum 3:1 ratio for protein precipitation, followed by vortex mixing for 15 seconds. Samples were centrifuged at 13,000 rpm for 7 minutes and the resulting supernatant was transferred to LC vials. The supernatant was stored at -80 °C until UHPLC-MS analysis, which was performed within one week. A blank sample, prepared with LC-MS grade water, underwent the same sample preparation process as the serum samples. Pooled quality control (QC) samples were prepared by combining 5-10 *μ*L aliquots of each serum sample extract. This pooled QC sample was analyzed every 10 LC-MS runs to monitor and correct instrument stability through the course of the experiment. Samples were randomized both for sample preparation and LC-MS analysis.

### Ultrahigh Performance Liquid Chromatography-Mass Spectrometry Serum Lipidomics

Reverse phase (RP) chromatography was performed in a Vanquish LC system equipped with a Thermo Accucore C30, 150 × 2.1 mm, 2.6 µm particle size column. An Orbitrap ID-X Tribrid mass spectrometer (ThermoFischer Scientific) was used for MS analysis. For negative ion mode, mobile phase A was 10 mM ammonium acetate with water/acetonitrile (40:60 v/v) and mobile phase B was 10 mM ammonium acetate with 2-isopropanol/acetonitrile (90:10 v/v). For positive ion mode, mobile phase A was 10 mM ammonium formate with water/acetonitrile (40:60 v/v) and 0.1% formic acid. Mobile phase B was 10 mM ammonium formate with 2-isopropanol/acetonitrile (90:10 v/v) and 0.1% formic acid. All samples were kept in the autosampler at 4 °C during LC-MS runs and an injection volume of 2 *µ*L was used in all cases. MS data were acquired in the 150-2000 *m/z* range with a 120,000 mass resolution setting. The most important MS parameters and the chromatographic gradient used are given in **Tables S2 and S3**, respectively. For MS/MS experiments, the Deep AcquireX data acquisition workflow was applied. Stepped normalized collision energy (NCE) values of 15, 30, 45 were used for fragmenting precursor ions in the HCD cell followed by Orbitrap analysis at 30,000 mass resolving power. Precursor ions were also fragmented with a CID energy of 40 and analyzed in the ion trap.

### Data Processing

Spectral features (retention time, *m/z*) pairs were extracted from raw data using Compound Discoverer v3.3 (Thermo Scientific). This step included chromatographic peak alignment, peak peaking, peak area integration, and adjustment for instrument drift using the pooled QC injections and the systemic error removal using random forest (SERRF) algorithm^29^. Chromatographic peaks with less than five times the peak area of the matching sample blank peaks were marked as background noise and removed from the dataset. Further filtering was performed by removing features not present in at least 50% of the QC sample injections or that had a relative standard deviation greater than 30% in QC samples. Following feature extraction, all retained features were matched against a curated in-house lipid spectral database. Exact masses, elemental formulas, and MS/MS spectra were used for matching purposes, and results manually curated. All annotated features were subject to ML feature selection.

### Quality Control

Data quality was assessed using the set of pooled QC runs. The average intensities of all QC runs were examined with RawMeat (Vast Scientific) software and their relative standard deviation (RSD) for positive and negative ion modes calculated. An average RSD below 15% was obtained for positive ion mode data. A slight drift in the negative ion mode dataset was observed, with a %RSD of 35%. For this dataset, features that could not be aligned after data processing in Compounds Discoverer were removed. Principal component analysis (PCA) showed excellent clustering of QC samples (**Figure S1**), confirming good reproducibility. Additionally, the quality of all sample and QC runs was evaluated across each batch using the IS peak areas (**Table S4**) and found to be excellent.

### Selection of Discriminant Lipids

Exploratory analysis of the overall lipidome alterations in OC was conducted using the following pipeline: first, one of a pair of two highly correlated features was removed using a Pearson’s correlation coefficient cutoff of 0.85. Fold changes for all remaining features were then calculated as the base 2 logarithm of the average lipid abundance ratios between OC and non-OC patients. Following this step, the statistical significance of each detected lipid was calculated using Welch’s t-test with a Benjamini Hochberg correction. Lipids with q-value < 0.05 were considered statistically significant. Altered lipid features were then autoscaled followed by feature selection using the SelectFromModel function with a random forest classifier in the Python sci-kit learn library (v1.1.2). Features were ranked by their Gini index; lipids with a Gini index equal to or greater than the Gini index mean were selected. The sci-kit learn default parameters were used and the number of trees for the random forest classifier was set to 100. To study differences between early-stage OC, non-OC and late-stage OC, the dataset was stratified by OC stages. The early-stage class consisted of stages I and II and the late-stage OC class was built using stages III and IV. Lipids statistically different between OC and non-OC groups (p-value < 0.05) were retained, followed by the random forests feature selection process described above.

### ML Pipeline for Biomarker Panel Selection

A small panel of lipids differentiating OC from non-OC samples was selected using the following ML pipeline: (1) One of two highly correlated features was removed using a Pearson’s correlation coefficient cutoff of 0.85. (2) The dataset was split into training (70%, OC *n* = 144 and non-OC *n* = 83) and test set (30%, OC *n* = 64 and non-OC *n* = 34). (2) Next, Welch’s t-test (p < 0.05) was applied to OC and non-OC samples in the training set (3) Since the dataset contained fewer number of non-OC samples than OC samples, the training and test sets were imbalanced and consisted of almost twice as many OC samples than non-OC. Imbalanced datasets often lead to poor classification performance as the classification classes are not equally represented^30^. To achieve improved classification power, the training set was balanced *via* the Synthetic Minority Over-sampling Technique (SMOTE)^30^, which can be used to create synthetic minority class samples for imbalanced datasets. Python’s imbalanced-learn library (v. 0.9.1) was used to implement SMOTE. (4) Feature selection was carried out on the balanced training set. In this case, random forests were used, and features were ranked by their Gini index feature important score. The top 10 lipids provided the best discriminating power for the training set and were selected for classification purposes. The sci-kit learn default parameters were used while the number of trees was set to 100. The classification power of ganglioside lipids was also evaluated. The best discriminating gangliosides were selected following the same feature selection process as described above. The top seven ganglioside features provided the best discriminating power for the training set and were selected for further analysis. Jupyter notebooks was used as the integrated development environment for all statistical and ML analysis. All code is available via GitHub: https://github.com/facundof2016/Sah_Ovarian-cancer_2023.

### ML Classification

Classification tasks were performed by training ML models to differentiate OC from non-OC samples. A balanced training dataset (*n* = 144 OC, *n* = 144 non-OC) was used to train the models, with a 10-fold cross validation conditions. Random forests, logistic regression, k-nearest neighbor (k-NN), linear kernel support vector machine (SVM-Lin) and a voting ensemble classifier were used. The estimators for the voting classifier included all four machine learning methods: random forests, logistic regression, k-NN, and SVM-Lin. The default parameters of Python’s sci-kit learn library (v1.1.2) were used. All classifiers were evaluated based on sensitivity, specificity, accuracy, and their area under the receiver operating characteristic curve (AUC ROC).

### AutoML Technique for Classification Tasks

For AutoML using auto-sklearn (version 0.15.0), the AutoSklearnClassifier was allocated 3600 seconds to identify the optimal ML pipelines. We utilized a cross-validation resampling strategy with standard parameters, adopting the hold-out method. This method further segmented the training data into an internal training and validation set with a 67:33 ratio. The ensemble approach was implemented, with consideration given to up to 50 models for inclusion in the ensemble. ROC AUC was used as the evaluation metrics. During optimization, each model was restricted to a maximum runtime of 1440 seconds. After establishing the AutoML pipelines using autosklearn, the final ensemble was trained on the complete training dataset through 5-fold cross-validation. Subsequently, its performance was assessed on unseen datasets.

## Results

### Patient Cohort

A high-coverage serum lipidomic LC-MS profiling workflow was applied to the combined patient cohorts. Each cohort was composed of ovarian cancer patients, benign ovarian tumors, benign uterine tumors, or cervical cancer patients. Patients with benign conditions and cervical cancers were grouped together as non-OC samples, while OC patients of various stages and histological types were all combined into the OC group. To remove potential confounders from the dataset, OC and non-OC groups were age matched. The statistical significance between the ages of OC and non-OC patients was assessed with Welch’s t-test. The age-matched cohort consisted of 208 OC and 117 non-OC patients **(Table 1 and Figure 1**). Among OC patients, 93 women had early-stage (I and II) cancers, of which 30% (28/93) were of serous histology, the most aggressive subtype^1^. In contrast, serous tumors accounted for 86% of advanced-stage (III and IV) cases while the remaining 14% of the cases included clear cell, transitional, mucinous, and carcinosarcoma subtypes.

**Figure 1.**
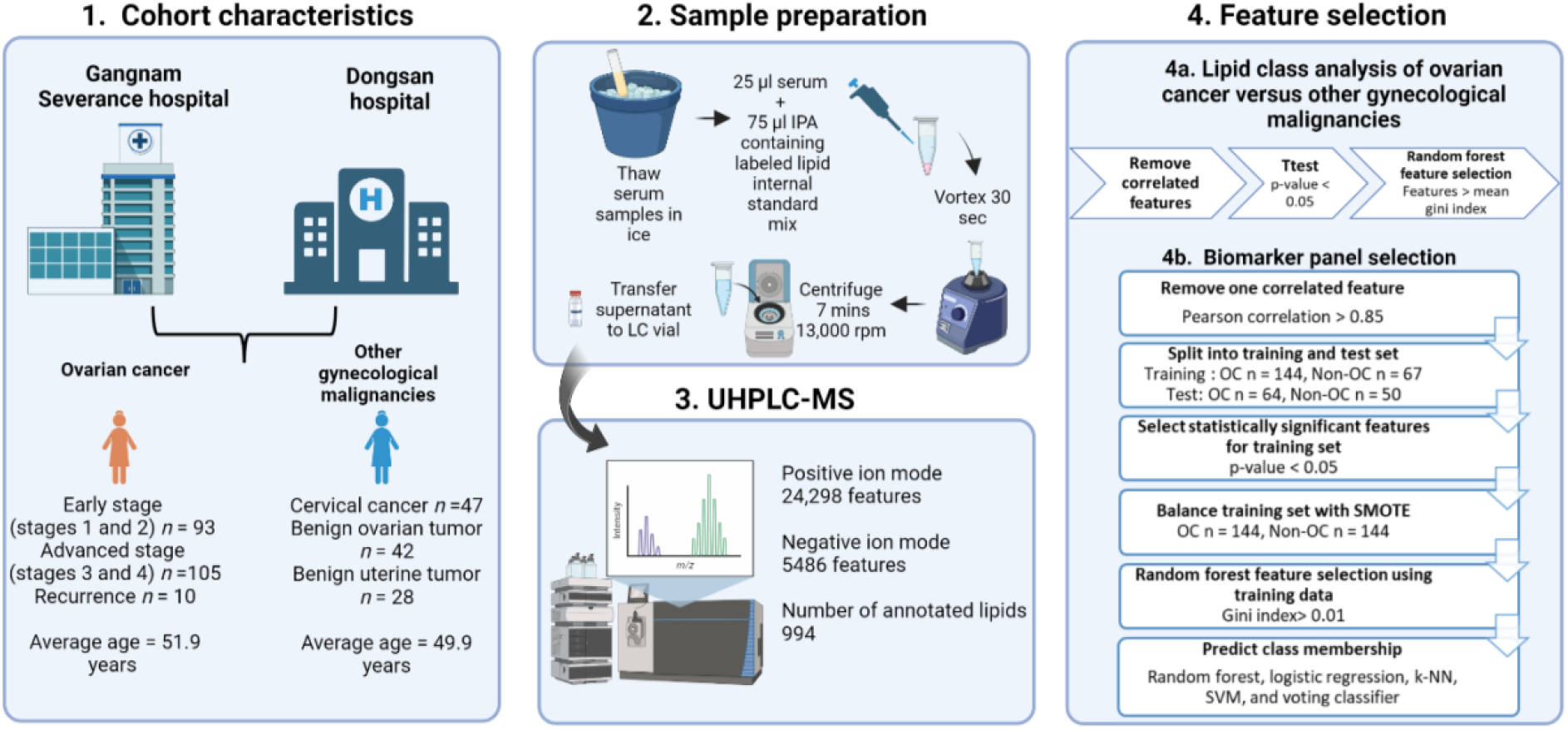
Study design overview, UHPLC-MS workflow, and machine learning pipeline. (1) Serum samples from OC (*n* = 208) and non-OC patients (*n* = 117) obtained from Gangnam Severance and Dongsan hospitals were studied. (2) Serum sample preparation workflow prior to UHPLC-MS analysis. (3) UHPLC-MS data collection in both positive and negative ion modes. Data were processed with Compound Discoverer v.3.3 (Thermo Scientific) and features annotated using in-house MS/MS libraries. (4) Machine learning workflow for selecting the most relevant lipids to differentiate OC and non-OC conditions. (4a) Selection of the best differential lipids using random forests feature selection. Lipids with a Gini index greater than the mean of all Gini index values were selected. (4b) Selection of a 10-lipid panel for differentiating OC and non-OC serum samples using random forests algorithm. Lipids with a Gini index of > 0.01 were selected as the best differential features in the training set. Five different machine learning models – random forests, logistic regression, k-Nearest Neighbor (KNN), support vector machines (SVM) and voting classifier, which is an ensemble of the four listed classifiers — were used for classification. SMOTE: Synthetic Minority Oversampling Technique. This figure was created in Biorender.com.

### Serum Lipidome Differences Between OC and non-OC patients

Lipidomics data were acquired for all OC and non-OC samples using reverse phase RP UHPLC-MS. A total of 24,297 and 5,485 de-isotoped and de-adducted spectral features (retention time, *m/z* pairs) were extracted from positive and negative ion mode datasets, respectively. From these, a total of 994 lipid species assigned to 22 lipid subclasses were successfully annotated using our in-house MS/MS spectral library. Detected lipid classes included fatty acids, glycerophospholipids, glycerolipids, sphingolipids, and sterol lipids. Triglycerides (TG) and phosphatidylcholines (PC) accounted for 29.6% and 18.2% of the dataset, respectively. Representative raw data showing separation of the various lipid classes detected are shown in supplementary information **Figure S2**, showcasing the excellent lipid coverage obtained in our method. Exploratory analysis to investigate lipidome differences among different OC histological types was first conducted. In addition to common lipids, 22 gangliosides were annotated by matching their elemental formulas and exact masses to online databases including LipidMaps^31^ and HMDB^32^. MS/MS information was available for 16 of the 22 annotated gangliosides and was also matched against online databases (**Table S5**). Principal component analysis (PCA) using the combined set annotated lipids did not show any clear clustering based on various OC histological types. (**Figure S3**). Unsurprisingly, the one sample with low-grade serous carcinoma (LGSC) histology deviated the most from the rest of the samples. This could be due to the low number of LGCS samples present in the dataset (*n*=1). Compared to high-grade serous carcinoma (HGSC), LGSC has a different mode of carcinogenesis with distinct molecular-genetic features^33^.

To investigate differences between OC and non-OC patients at the lipidome level, unsupervised and supervised multivariate analysis was conducted. PCA using the combined set of (+) and (-) RP UHPLC-MS 994 annotated lipids showed minimal separation between OC and non-OC patients (**Figure S4A**). This was expected as differences between different types of malignancies are likely to be relatively subtle. The dataset was further explored with supervised multivariate analysis. Orthogonal partial least squares-discriminate analysis (oPLS-DA) for the same dataset indicated better clustering between the two groups (**Figure S4B**), which could further be improved *via* ML-based feature selection processes. Performance characteristics of this oPLS-DA model with 10-fold cross validation were a modest 0.73 sensitivity and 0.65 specificity. To better visualize lipidome differences between OC and non-OC patients, the dataset was analyzed on a lipid class basis. Fold changes and statistical significance between OC and non-OC groups were calculated for all annotated lipids (**Figure 2**). Among the 994 annotated lipids, 218 had false-discovery rate (FDR) corrected p-values < 0.05 (q< 0.05). Fold changes calculated between OC and non-OC groups showed an overall decrease in the serum lipid abundances of OC patients. Seventy-four percent of the detected ether phosphatidylethanolamines (PE O-), for example, had significantly decreased abundance in OC patients. Lipid classes including PC, ether phosphatidylcholines (PC O-), lysophosphatidylcholines (LPC), and lysophosphatidylethanolamines (LPE) were also significantly reduced in women with OC (**Figure 2**). Although most lipids were reduced in the OC samples, some lipids including ceramides (Cer) and TG were increased. Interestingly, the only two ceramides species that were lower in OC patients were Cer(d18:0/22:0) and its oxidated form Cer(t18:0/22:0).

**Figure 2.**
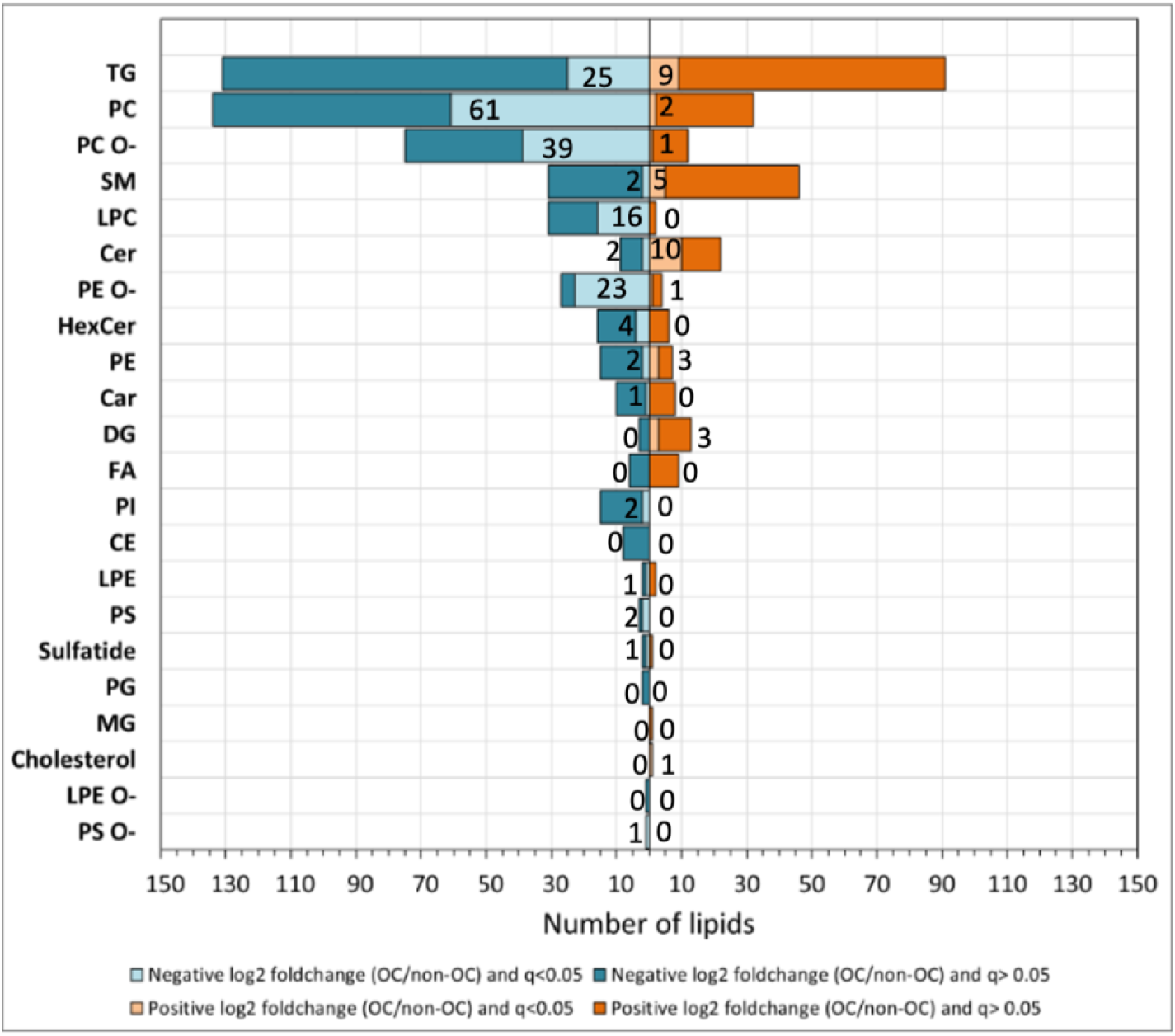
Annotated lipids, grouped by lipid class, showing number of lipids with positive and negative fold changes between OC and non-OC groups. Fold changes were calculated as the base 2 logarithm of the average lipid abundance ratios for OC *vs.* non-OC. A positive fold change value indicates higher levels in OC samples. Negative values indicate lower levels in OC samples. The number of lipids with negative fold change values and positive fold change values for each lipid class are shown as blue and orange bars, respectively. The number of statistically significant lipids (FDR corrected p-value < 0.05) with negative and positive fold change values are labeled as light blue and orange bars, respectively. TG: Triacylglycerols, PC: Phosphatidylcholines, PC O-: Ether phosphatidylcholines, SM: Sphingomyelins, LPC: Lysophosphatidylcholines, Cer: Ceramides, PE O-: Ether phosphatidylethanolamines, Car: Carnitines, HexCer: Hexosylceramides, PE: Phosphatidylethanolamines, DG: Diacylglycerols, FA: Fatty acids, PI: Phosphatidylinositols, CE: Cholesterol esters, LPE: Lysophosphatidylethanolamines, PS: Phosphatidylserines, PG: Phosphatidylglycerols, MG: Monoradylglycerols, LPE O-: Ether Lysophosphatidylethanolamines, PS O- : Ether phosphatidylserines.

A ML workflow, described in the methods section and **Figure 1**, was developed to select features that better describe lipidome differences between OC and non-OC conditions. The one hundred best differential lipids were selected (**Figure 3**), including ceramides, LPC, PC, PC O-, PE O-, and TG, indicating diverse changes in the circulating lipid abundances in serum of OC patients. Unsupervised PCA and supervised oPLS-DA using these selected 100 lipids (**Figure 3**) showed better clustering than the previously built models using all annotated lipids (**Figure S4**). Performance characteristics of the oPLS-DA model on the first 2 latent variables were 0.75 and 0.71 cross-validated sensitivity and specificity, respectively. Comparison of the relative lipid abundances in OC and non-OC conditions showed patterns of alterations based on the lipid class, with majority of the lipids showing decreased serum abundance in OC patients. Only ceramides, some SM, cholesterol, FA(20:1), diglycerides, monoglyceride (MG), and some TG species were increased in the OC patients. TG species showed variable trends based on the FA side chain: most TG species with long fatty acyl side chain composition were increased while TG with short fatty acyl side chains were decreased. In addition, fifteen out of the 100 selected lipids had significantly different fold changes between OC and non-OC groups (log_2_ fold change < -0.5 or log_2_ fold change > 0.5), confirming lipids exhibit profound changes in ovarian cancer. These mainly consisted of PE O-, PC O-, and PC.

**Figure 3.**
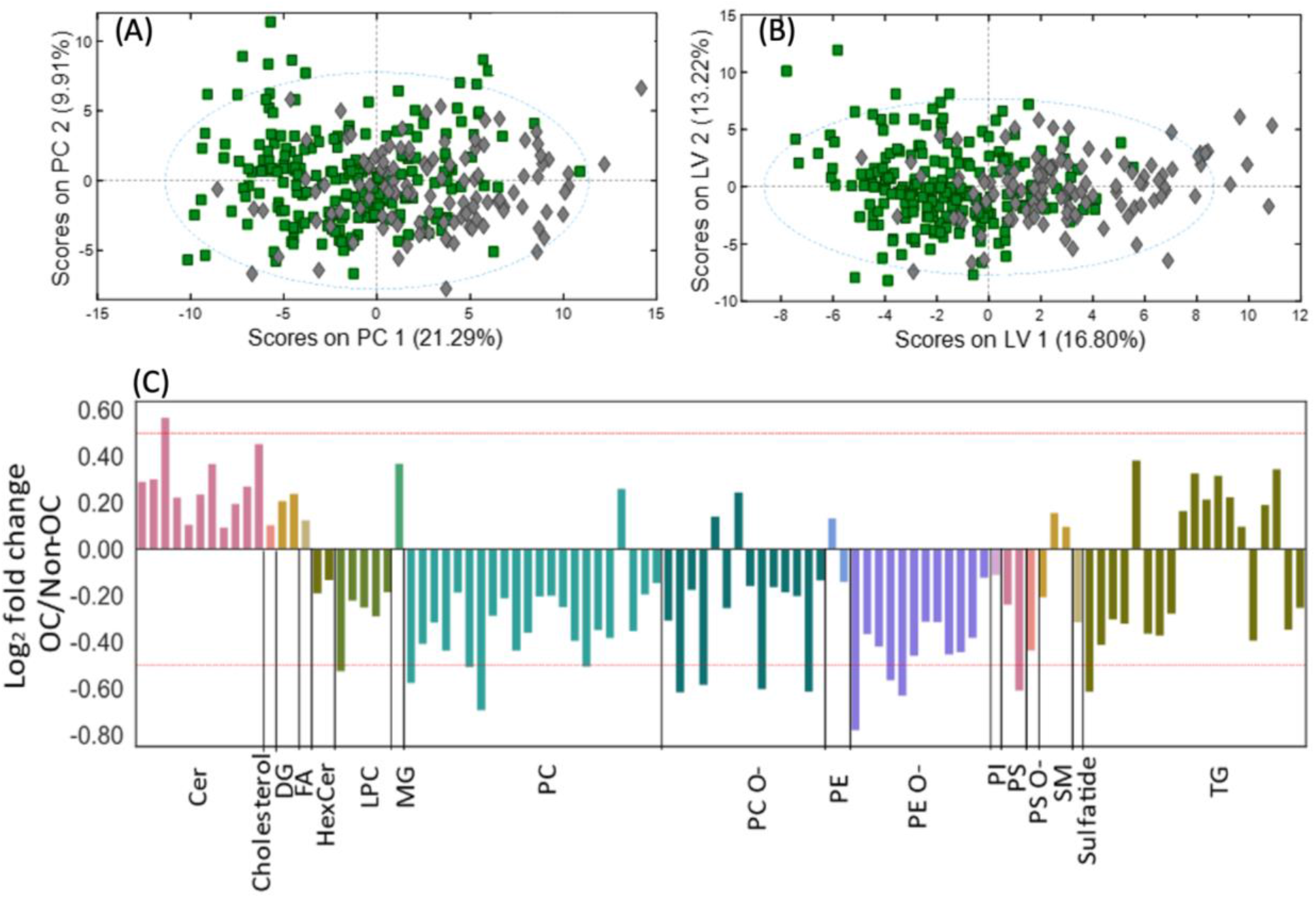
Serum lipidome analysis of OC and non-OC samples using only the abundances of select lipids. Lipids were selected with the following feature selection workflow: one of the two highly correlated lipids was filtered out using a Pearson correlation coefficient cutoff of 0.85. Next, lipids with *p*-values lower than 0.05 were selected, followed by random forest feature selection in which lipids with a Gini index greater than the mean of all Gini indices were selected. (A) PCA score plot showing clustering of OC and non-OC samples using the selected lipids. (B) o-PLS-DA score plot for the same dataset. OC samples are depicted as green squares and non-OC samples are shown with grey diamonds. (C) Fold changes for the selected lipids. Fold changes were calculated as the base 2 logarithm of the average lipid abundance ratios for OC *vs.* non-OC. A positive fold change value indicates higher levels in OC samples. Negative values indicate lower levels in OC samples. TG: Triacylglycerols, PC: Phosphatidylcholines, PC O-: Ether phosphatidylcholines, SM: Sphingomyelins, LPC: Lysophosphatidylcholines, Cer: Ceramides, PE O-: Ether phosphatidylethanolamines, Car: Carnitines, HexCer: Hexosylceramides, PE: Phosphatidylethanolamines, DG: Diacylglycerols, FA: Fatty acids, PI: Phosphatidylinositols, CE: Cholesterol esters, LPE: Lysophosphatidylethanolamines, PS: Phosphatidylserines, PG: Phosphatidylglycerols, MG: Monoradylglycerols, LPE O-: Ether Lysophosphatidylethanolamines, PS O- : Ether phosphatidylserines.

### Lipidome Analysis of High-Grade Serous Carcinomas

The heterogeneity of OC is one major obstacle in biomarker discovery^23^. HGSC accounts for only 30% of early-stage OC, but 86% of advanced-stage OC. The overwhelming majority of early-stage OCs (∼69%) are non-HGSCs, the OC types that are generally slow-growing, less aggressive and therefore commonly detected at early stages^1^. In contrast, most advanced-stage OCs (86%) are HGSC, which is aggressive and typically not detected until advanced stage. These findings indicate that, potentially, HGSC has biological mechanisms of development and progression distinct from non-HGSCs. We investigated whether HGSC exhibits a distinctive lipid profile compared to other OC subtypes. Welch’s t-test was performed to select statistically different features between HGSC and other OC types. Seventy-four lipids were selected as statistically significant (p< 0.05) however when p-values were corrected for false positives (FDR corrected) no lipids remained statistically significant, indicating lipidome differences between different histological types are likely small, or not present in the subset of annotated lipids. Nevertheless, we still examined the lipidome of HGSC with the 74 selected features with p-value < 0.05. Comparison of the serum lipid abundance of HGSC *vs.* non-HGSC OC types showed subtle differences (**Figure S5**). Only one TG species showed a log_2_ fold change greater than 0.5. Overall, the serum lipidome of HGSC showed increased lipid abundance compared to non-HGSC types. Most lipid classes including all ceramides, and most glycerophospholipids, and glycerolipids were increased in HGSC, while only ether phospholipids including PE O- and PC O- and a few SM, PC and PE species were reduced. To further investigate the serum lipid profile of HGSC, we compared the serum lipidome of HGSC with non-OC malignancies. Welch’s t-test selected 202 statistically significant features with FDR corrected p-values < 0.05. Fold changes between HGSC and non-OC showed that, unlike HGSC *vs.* other OC subtypes, in this case most lipid classes were reduced in HGSC (**Figure S6**). These results bear similarities with our previous analysis of all OC subtypes *vs.* non-OC (**Figure 3**), reinforcing the finding that lipid alterations between OC and non-OC conditions are much more profound than they are between different histological OC types.

### Stage Stratified Analysis

Next, we evaluated the lipidome profile associated with early-stage OC (I and II) and advanced-stage OC (III and IV) versus non-OC conditions using a stage-stratified feature selection approach, as described in the methods section. The early-stage OC cohort consisted of 93 OC patients and 117 non-OC patients, while the late-stage OC cohort had 115 OC and 117 non-OC patients. One hundred twenty lipids were selected as the best differential lipids between early OC and non-OC while 102 lipids were selected for late stage *vs.* non-OC conditions. While fewer lipids were selected for late-stage OC *vs.* non-OC than early-stage OC *vs.* non-OC, a greater diversity of lipid classes was altered for late-stage OC (**Table S6 and S7**). Both early and late-stage OC exhibited changes in glycerophospholipids, glycerolipids, and sphingolipids, while some lipid classes—including diglycerides, fatty acids, and cholesterol—differed only for late-stage OC patients. Most of the selected differential lipids for both early-stage and late-stage OC included PC, TG, PC O-, PE O-, and Cer (**Figure 4**). Compared to late-stage OC, a higher number of LPC and SM species were selected for early-stage OC. Furthermore, as expected, lipidome alterations in early disease stages were less prominent than in advanced disease stage. Volcano plots showing the fold changes and statistical significance for the selected lipids show that ceramides, some TG, and SM were increased in both early and late-stage OC. Lipid classes identified as significantly altered (p-value < 0.05 and fold change greater than log_2_ 0.5) in early-stage OC included PE O-, PC O-, PC, and one PS specie (**Table S8**). Late-stage OC exhibited significant changes in Cer, PE O-, PC O-, PC, LPC, PS, and TG **(Table S8**).

**Figure 4.**
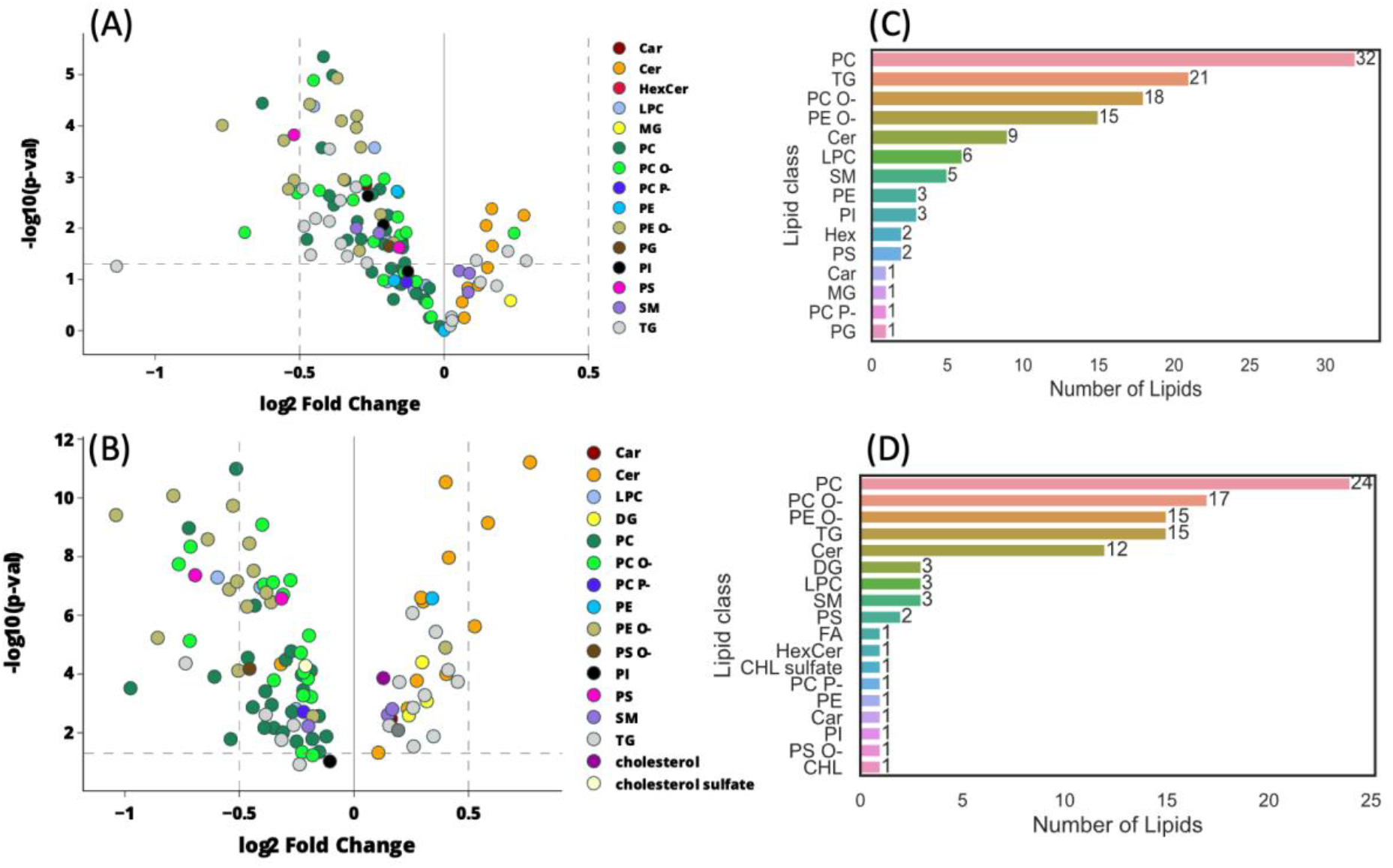
Serum lipidome differences in early stage (I and II) or advanced stage (III and IV) OC *vs.* non-OC patients using only select lipids. Most relevant lipid species for differentiating early-stage OC from non-OC, and advanced stage OC from non-OC were selected using random forests feature selection. Volcano plots showing lipidome differences with (A) 120 best discriminating lipids selected for early-stage OC *vs.* non-OC (B) 102 best discriminating lipids selected for advanced stage OC *vs.* non-OC samples. Lipid species are color-coded by lipid class, as indicated on the plots. Fold changes were calculated as the base 2 logarithm of the average lipid abundance ratios for OC *vs.* non-OC. A positive fold change value indicates higher levels in OC samples. Negative values indicate lower levels in OC samples. *p-*values were calculated using Welch’s t-test. Bar graphs showing number of lipids selected, grouped by lipid class, for (C) early-stage OC *vs.* non-OC (D) advanced stage OC *vs.* non-OC. TG: Triacylglycerols, PC: Phosphatidylcholines, PC O-: Ether phosphatidylcholines, SM: Sphingomyelins, LPC: Lysophosphatidylcholines, Cer: Ceramides, PE O-: Ether phosphatidylethanolamines, Car: Carnitines, HexCer: Hexosylceramides, PE: Phosphatidylethanolamines, DG: Diacylglycerols, FA: Fatty acids, PI: Phosphatidylinositols, CE: Cholesterol esters, LPE: Lysophosphatidylethanolamines, PS: Phosphatidylserines, PG: Phosphatidylglycerols, MG: Monoradylglycerols, LPE O-: Ether Lysophosphatidylethanolamines, PS O- : Ether phosphatidylserines, CHL: cholesterol.

Among the differentially abundant lipids selected for early-stage or late-stage OC *vs.* non-OC conditions, 42 lipids were altered in both subgroups (**Figure S7**). These were mainly sphingolipids including Cer and SM, glycerophospholipids including PC, PS and PI, ether phospholipids (PC O- and PE O-) and TG species, Lipids with fatty acid alkyl chains of C18:0, C18:1, C18:2, C20:4, C22:5, and C22:6 were frequently observed in this panel. In addition, all selected ceramide species were composed of d18:1 or d18:2 fatty acyl backbone, while the fatty acyl side chains were very long fatty acids, including C24, C25, or C26. Similarly, the sn-1 fatty acid alkyl chain for most PE O-, PE, and PS species were composed of C18:1 or C18:2, whereas sn-2 fatty acid alkyl chains were composed of very long chain fatty acyl chains of C22 and C24. This panel of lipids also consisted of glycerophospholipids with dietary odd fatty acyl chain composition (C17:0, C17:1, and C17:2). Next, fold changes for these 42 lipids between early- or late-stage OC *vs.* non-OC patients were analyzed. Subtle changes in serum lipid abundance were observed for early-stage OC *vs.* non-OC. Among the selected lipids, PE O-(17:1/20:4) had the greatest fold change. Lipid alterations for advanced stage OC *vs.* non-OC were much more profound than in early-stage OC patients, and 42 lipids showed alterations that were directionally concordant with the disease stage. These results indicate that changes in the serum lipid abundance of OC can be detected when the cancer is localized, and that these changes are amplified as the diseases progresses (**Figure S7**).

Additionally, we evaluated the serum lipidome of early-stage HGSC samples using the 120 lipids selected as differential features for early-stage OC *vs.* non-OC (**Figure S8**). In line with our lipidome analysis of HGSC shown earlier (**Figure S6**), the serum lipidome of early-stage serous patients exhibited alterations that in most cases were like the combined early-stage OC cohort of various histological types (**Figure 4**). Only a few PC species showed increased abundance in early-stage serous samples. In addition, five lipids, including TG(56:6), PC(O-35:4), PC(O-32:2), PE(O-18:2/20:1), and PC(37:3), showed significant alteration (**Figure S8**), suggesting that circulating lipids can aid in early diagnosis of serous carcinoma.

### Biomarker Panel

Considering the importance of biomarkers in OC diagnosis and management, we selected a panel of lipids to discriminate OC and from non-OC conditions with the maximum possible accuracy. All 994 annotated lipids from (+) and (-) RP UHPLC-MS datasets were combined and subjected to the ML workflow detailed in the methods section and **Figure 1**. Although the AUC values for the imbalanced dataset (OC *n* = 144, non-OC *n* = 83) were acceptable, the specificity for these classification models were very low (**Table S9**). For the imbalanced dataset a logistic regression model provided the best AUC value of 0.76 while the specificity for this model was only 0.50. Thus, to achieve better classification power the training set was balanced using the Synthetic Minority Oversampling Technique (SMOTE)^30^, yielding a 10-lipid panel that consisted mainly of ceramides and ether linked glycerophospholipids (**Table 2**). The best classification performance for the training set was achieved with a random forest classier: the AUC, sensitivity, and specificity under 10-fold cross validation were 0.85, 0.78, and 0.76, respectively (**Table S10, Table 3**). For the test set, the random forest classifier provided the AUC of 0.82 while the sensitivity and specificity values were 0.78 and 0.76, respectively. The selected panel of lipids provided good classification performance for the test set, highlighting the potential of lipid markers to distinguish the serum lipid profile of Korean women with OC and different types of malignant or benign gynecological diseases.

**Table 2.**
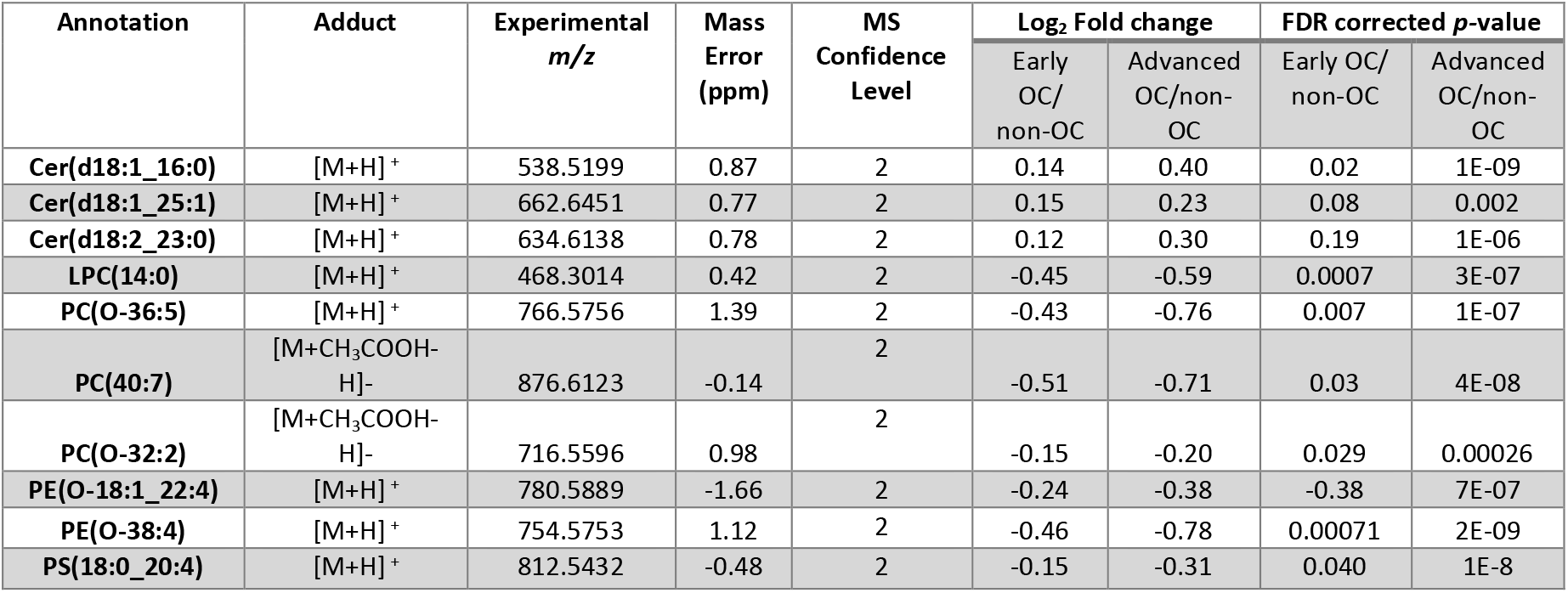
Discriminant (OC *vs.* non-OC) ten-lipid panel selected using a random forests algorithm. Proposed lipid annotation, main adduct type detected, experimental monoisotopic *m/z* value, mass error (ppm), MS annotation confidence level, *p*-values, and abundance log-transformed fold changes are shown. MS annotation level was assigned based on the following criteria: 1) MS1 and MS/MS spectrum of standard matched to the feature. 2) MS1 and MS/MS spectrum of the feature matched with library spectra 3) tentative ID assignment based on elemental formula match with literature. 4) unknowns.

**Table 3.**
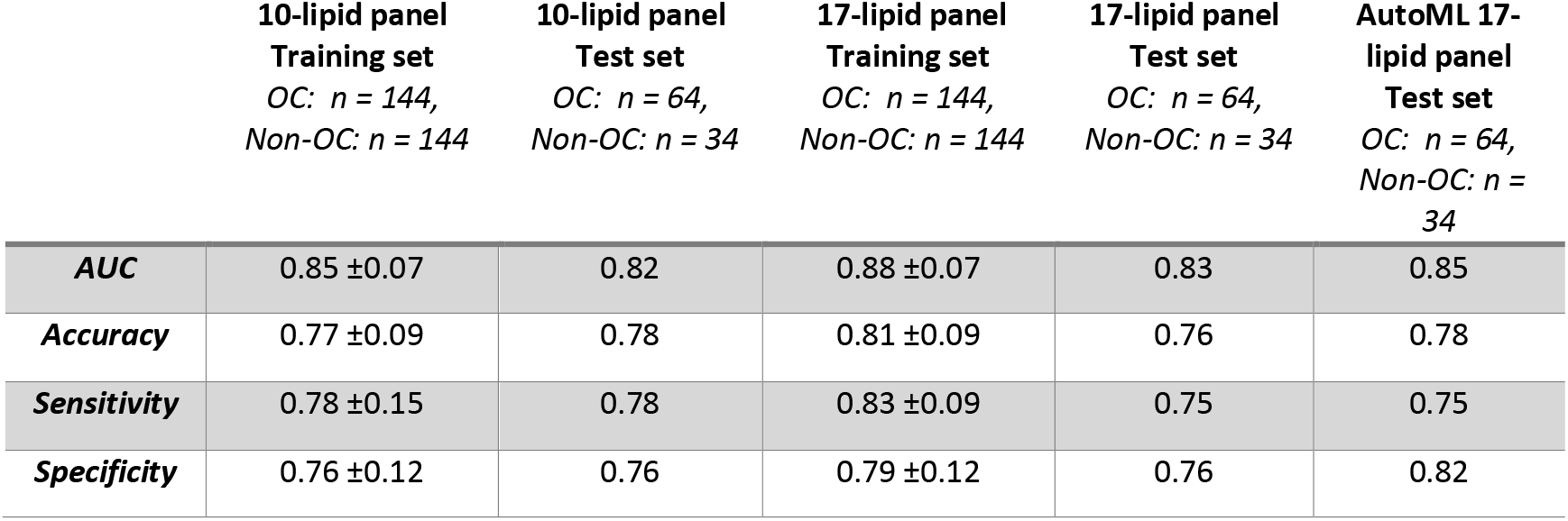
Machine learning performance of various lipid panels. Classification performance using a random forest classier and the AutoML model. The top 7 discriminant gangliosides were added to the 10-lipid panel to create the 17-lipid biomarker panel. Training data were balanced with the Synthetic Minority Oversampling Technique (SMOTE).

As accurate early diagnosis of OC, especially of early-stage HGSC is crucial to improve clinical outcomes, we evaluated the classification power of this 10-lipid panel for early-stage samples (*n* = 31) *vs.* non-OC samples (*n* = 30) from the test set, as well as for all 28 early-stage HGSC samples (samples used in training set and test set) *vs.* non-OC samples (*n* = 30). Random forest classification model for early stage *vs.* non-OC samples gave the AUC of 0.75 while the specificity was only 0.45 (**Table S10**). Better specificity could be achieved by selecting a panel of lipids specifically for discriminating early-stage OC from non-OC conditions. Additionally, since the non-OC dataset consisted of invasive cervical cancer samples it is not surprising to obtain low classification performance as we are discriminating early-stage OC samples from other cancers. Improved classification performance was achieved for a sample set without cervical cancer samples (**Table S11**). The 10-lipid biomarker panel discriminated early-stage OC from benign conditions (without cervical cancer) with the AUC, sensitivity, and specificity of 0.86, 0.81, and 0.69, respectively.

Comparison of the relative abundances of these 10 selected lipid markers between early-stage OC, late-OC, and non-OC patients showed trends following the course of the disease (**Figure 5**). Ceramides showed an increasing trend while phospholipids decreased with the disease progression. Although we were not able to compare diseased patients against normal controls, the directionally concordant changes in lipid abundances suggest the lipidome profile of women with gynecological malignancies can be monitored for improved clinical triage.

**Figure 5.**
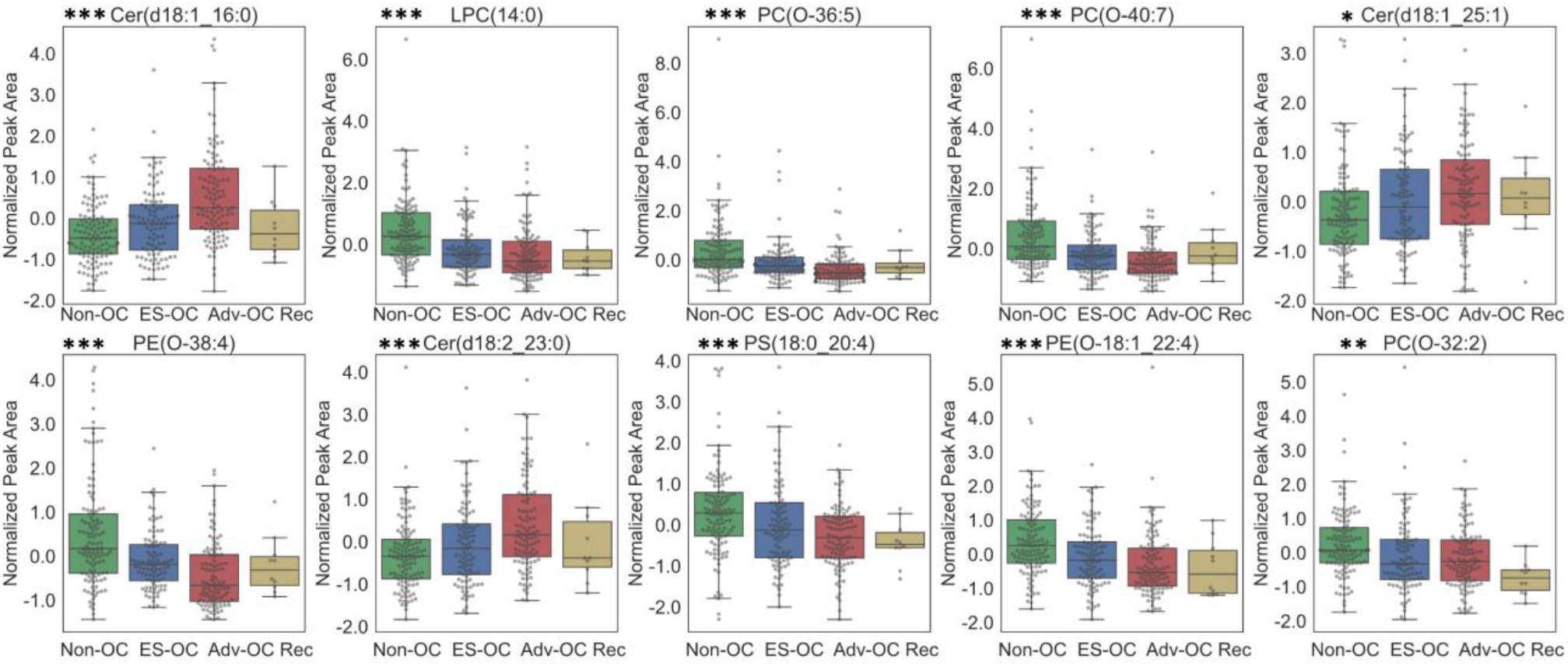
Relative abundances for the ten discriminant lipids in early OC, advanced OC, and non-OC samples. *p*-values between OC and non-OC samples were calculated using Welch’s t-test. (* p < 0.05, ** p ≤ 0.01, *** p ≤ 0.001). Raw data were autoscaled for visualization. ES-OC: early-stage ovarian cancer, Adv-OC: advanced stage ovarian cancer, Rec: recurrent OC.

### Serum ganglioside alterations with OC

Previous studies have indicated increased gangliosides levels in OC cell lines, tissues, serum, as well as ascites fluid from OC patients^23, 24^, suggesting gangliosides could play a role as OC markers. However, existing data is still sparse regarding the importance of these glycolipids in OC development^23^. A total of twenty-two ganglioside species were detected in serum by LC-MS (**Table S5**), and their structures confirmed by tandem MS experiments. Examination of the average ganglioside abundances for early-stage OC, advanced-stage OC, and non-OC patients showed an increase in overall ganglioside abundances concurrent with disease progression (**Figure S9**). However, the changes in ganglioside abundances between non-OC and OC patients were not significant if tested univariately. Therefore, the diagnostic power of gangliosides was explored by combining them in panels. Classification models were built using all annotated gangliosides, which showed acceptable performance. The best random forest model provided AUC values for training and test sets of 0.78 and 0.68, respectively. To investigate whether gangliosides could further enhance the diagnostic power of the previously selected 10-lipid panel (**Table S10, Table 2**), the top 7 discriminating gangliosides features in these models were added to the panel, resulting in a new 17-lipid panel (**Table S12**). Improved AUC values were obtained for training and test sets with this set of lipids: the best AUC values for the training and test set were 0.88 and 0.83, respectively (**Table S13 and Table 3**). As with previous comparisons, the training set was balanced using SMOTE and was cross-validated 10-fold.

### AutoML Classification

In addition to traditional ML approaches an AutoML method, Auto-sklearn^28^, that automatically identifies optimal ML pipelines for a given task, was applied. The Auto-sklearn technique employs Bayesian optimization, meta-learning, and ensemble techniques to automatically select ML pipelines. Through Bayesian optimization, it assesses hyperparameter settings for optimal results, while meta-learning helps to learn from past experiences to enhance performance on new tasks. The framework also constructs ensembles by selecting models that minimize training data errors. Here, the AutoML technique (detailed in methods section) provided improved AUC, accuracy, and specificity for the test set using the 17-lipid biomarker panel. Classification metrics using auto-sklearn were 0.85, 0.78, 0.75 and 0.82 AUC, accuracy, sensitivity, and specificity, respectively (**Table 3**).

### Serum Lipidomics of Normal Controls vs. OC Patients

Blood samples from normal controls were collected for 150 women during regular checkups. Although these controls were excluded from the feature selection process, we investigated whether lipids selected for differentiating OC from non-OC patients could also distinguish normal controls. Clustering between OC and normal controls was investigated using the full set of annotated lipids (**Figure S10**). Fold changes between OC and normal controls for these 100 select lipids also showed lower serum lipidome abundances in OC patients (**Figure S11**). Ceramides and TG showed variable trends with most ceramides showing increased abundances in OC patients. Compared to our previous analysis of OC *vs.* non-OC conditions (**Figure 3**), lipid alterations between OC and normal controls were much more profound. Several lipid species showed significant differences: PC(18:0/22:6) and LPC(18:2) were reduced approximately 10- and 6-fold in OC patients, respectively. The classification performance of the selected 10-lipid maker panel was also evaluated for age-matched OC *vs.* normal controls (**Table S10**). Since the dataset had a larger number of OC samples than normal controls, SMOTE^30^ was applied to balance the number of samples. Classification performance for the balanced set of OC patients *vs.* normal controls was evaluated using five ML methods: random forest, SVM, logistic regression, k-NN, and voting method. The best results were obtained with a random forest model. Performance characteristics of this model were 0.94, 0.88, 0.91, and 0.84 for the AUC, accuracy, sensitivity, and specificity, respectively.

## Discussion

Accurate distinction between ovarian cancer and other benign or cancerous gynecological malignancies remains an unmet clinical challenge with significant impact on patient survival^7^. Significantly improved survival rates are observed when women with OC are referred to tertiary care specialists rather than general gynecologists, and although treatment of women with benign diseases by specialists imposes no harm, benign disease misdiagnosed for cancer causes unnecessary burden and patient anxiety^6, 34–37^.

In recent years, dysregulation in lipid metabolism has been established as a crucial feature of ovarian cancer progression, reflecting the increased energy demands of highly proliferating cancer cells and cell membrane remodeling^38^. However, previous ovarian cancer lipidomic studies have been limited by low numbers of early-stage patients, low lipidome coverage, and biases originating in the sample collection process^16, 17, 22^. Although alteration in lipid metabolism is a characteristic feature of OC, findings from different studies still show numerous inconsistencies^38^. Previous data show racial/ethnic differences in risk, incidence, and survival of OC patients^39–41^, yet most studies consist mainly of non-Hispanic white women with European ancestry. Compared to other racial groups, Asian women have a higher incidence rate of mucinous ovarian tumors^42^ and it has been shown that the protein biomarker HE4 may be more useful for that cancer in this ethnic group^41^. It is generally agreed that studies in specific racial/ethnic groups are needed to better understand OC diversity and accelerate preventive and diagnostic strategies.

The study presented here presents results for the comprehensive analysis of the serum lipidome of OC patients of Korean descent, recruited from two different clinical sites. A consistent decrease in the overall lipid abundance across various OC stages and histological types was observed in our dataset when compared against non-OC conditions. These results agree with previous studies showing an overall abundance decrease in serum glycerophospholipid abundances in OC patients of non-Hispanic white and European decent^16, 17, 22^. This overall decrease in lipid abundance is likely due to altered lipoprotein level, as suggested in previous studiess^43, 44^. High-density lipoprotein (HDL) particles are rich in phospholipids; decreased HDL-cholesterol levels have been reported in OC patients^43^. Further, stage-stratified analysis comparing early- or advanced-stage OC *vs.* non-OC (**Figure 4 and Figure S7**) suggests that lipid alterations are detectable even in the early stages, amplifying as the disease progresses. These results further underscore the potential role of circulating lipids in monitoring OC development and progression. Similar results were reported by Buas *et al.* when comparing benign diseases with OC^22^ and recent studies suggest changes in circulating lipid abundances in pre-diagnostic specimens^45^, indicating lipid profiles exhibit changes years before OC diagnosis. Mounting evidence suggests that monitoring lipid profiles could aid in identifying women who are at higher risk of developing OC.

Although most lipids exhibited wide-scale abundance reductions in OC patients, some lipid classes including Cer, some TG species, and DG, showed a consistent increase across the disease spectrum. Increase in blood ceramides levels have been frequently reported for OC^17, 21, 46, 47^, with significant increase in specific ceramides including C16, C20, C22, and C24 derivatives. In our study, a consistent increase in almost all statistically significant ceramide species was observed for OC patients. Ceramides identified as differentially abundant for OC included those with d18:0, d18:1 and d18:2 fatty acyl backbones with C16, C23, C24, C25, and C26 fatty acid chains. Several studies have noted distinctive Cer alterations based on fatty acid chain length profiling. Studies have suggested an increase in ceramides with 16:0, 18:0, 20:1 and 24:1 fatty acid chain, while decreased abundance of 23:0 and 24:0 fatty acid Cer has also been reported. In our study, however, no such trend was observed, and all key Cer species showed increased abundance in OC. Although understanding the biological basis of these lipid alterations in OC is an active area of investigation, literature suggests increased Cer levels may be associated with the increased activity of ceramide synthases that promote tumor growth^48^. The frequently observed dysregulation of ceramides in OC along with their biological importance relative to tumor growth calls for a deeper understanding of OC sphingolipid metabolism.

Along with increased Cer levels, reduced abundance of PE O- and PC O- is also frequently reported in OC studies^16^. Here, we observed 74% of the detected PE O-species to be significantly decreased in OC patients, suggesting that PE O- are of particular importance in OC pathogenesis. Additionally, ratios between Cer and ether phospholipids abundances have frequently been reported as potential markers for OC^47, 49^, suggesting an important association between sphingolipid and ether-phospholipid metabolism^50^. Four out of the best 10 discriminating lipids were ether phospholipids and 3 were ceramides (**Table 2**) highlighting their diagnostic value and reinforcing the notion that a further understanding of the interplay between these lipid families in much needed in the context of OC.

A recent study has suggested gangliosides GD2 and GD3 as potential diagnostic OC^23^. Gangliosides are a class of glycolipids present in the plasma membranes of almost all vertebrate cells and are crucial in biological processes that include cell recognition and adhesion, transmembrane signaling, cell growth and differentiation^51^. Gangliosides are known to be involved in tumor-host immune system interactions^52^, and studies have reported pronounced local defects in OC immune responses^53, 54^. Gangliosides are shed into the extracellular environment, particularly during the malignant transformation of cells^55, 56^, making them ideal biomarker candidates. Past studies have reported lower survival rates of cancer patients with increased circulating ganglioside levels^24^. Despite the potentially important role of gangliosides in OC, however, reports on ganglioside expression in OC patients are quite sparse. In our study, we conducted in depth profiling of gangliosides, qualitatively observing an increased average abundance in OC patients compared to non-OC conditions. Abundance differences, however, were not statistically significant. A generally increasing trend in abundance with disease malignancy was observed, potentially reflecting increases in ganglioside secretion associated with tumor development^24^. These results are in line with previous studies reporting significantly higher ganglioside levels in OC cells compared with non-OC malignancies^24^. Additionally, our lipid panel results showed a moderate AUC improvement with the addition of 7 select gangliosides (**Table S13, Table 3**). Further improvement was achieved with the use of the developed AutoML method (**Table 3**). Future quantitative studies with enhanced sensitivity and specificity for gangliosides are likely to facilitate biomarker discovery and solidify the development of therapeutic targets for OC.

One salient characteristic of our study was the relatively large sample size for OC patients *(n* = 208), including 93 patients with early-stage OC (stage I and II). Securing samples from early-stage OC patients is a major challenge and thus most studies reported only on a very limited number of early-stage cases. In addition, as mentioned previously, most studies are limited to samples from non-Hispanic white women. Here, a stage-stratified analysis for 93 early-stage OC samples *vs.* 117 non-OC samples selected a variety of lipid classes as differential lipids (**Figure 4 and Figure S7**). Among the selected lipids, ether glycerophospholipids exhibited significant changes in early disease stage emphasizing their potential for distinguishing OC from other gynecological malignancies when the cancer is localized. Of all early-stage samples, the cohort included 23 patients with early stage HGSC – the deadliest subtype of OC^57^. Only 13% of serous ovarian carcinomas are diagnosed when this cancer is localized, leading to poor prognosis and very low survival rates. Our lipidome analysis of early-stage patients of serous histology showed systematic alterations in lipid abundance based on lipid class. Although, most lipid alterations for early-stage serous and early-stage OC samples of various histological types showed the same overall trend, an increased abundance of some specific PC species was observed only in the early-stage serous subtype (**Figure 4 and S8**), indicating some lipid alterations may be unique to HGSC. Alterations in the majority of lipid classes, however, followed a pattern similar to the other histological types. As indicated in **Figure S5**, as well as the exploratory PCA analysis of different OC subtypes (**Figure S3**), very subtle differences were observed between different OC histological types. Meanwhile lipidome analysis between HGSC *vs.* non-OC conditions (**Figure S6**) and all OC subtypes *vs.* non-OC (**Figure 3 and 4**) exhibited more prominent changes, suggesting changes in lipid profile associated with OC *vs.* non-OC cases surpass most of the differences caused by the histological diversity of OC. These findings underscore the daunting challenge of identifying OC subtype-specific biomarkers. Our results are in line with previous metabolomics studies^16^, but further analysis with larger samples of especially of early-stage serous carcinoma patients will be required to confirm these observations as well as to identify makers unique to HGSC.

The dataset presented here also provides an opportunity to contrast lipid changes associated with OC against cervical cancer. Data suggests that the OC lipidome profile is distinguishable from that of cervical cancer, opening the possibility of developing screening tools to simultaneously test for both ovarian and cervical cancers. Since testing for cervical cancer *via* Pap tests has been routinely performed for over 50 years, developing a lipid-based Pap tests screening strategy for both cervical and ovarian cancer could lead to improved OC diagnosis and disease prognosis^13^. In the future, we expect that further diagnostic gains could be achieved by integrating lipid/glycolipid markers with existing protein biomarkers such as CA125 and/or HE4^22^.

## Conclusions

In-depth metabolomic analysis of the serum lipidome of several benign and malignant gynecological diseases in Korean women revealed lipids unique to ovarian cancer. In comparison to benign ovarian and uterine tumors and invasive cervical cancer, the serum lipidome of ovarian cancer shows reduced abundance of most lipid classes, except for a few specific cases. Although OC metabolomics still show some level of disagreement, especially regarding PC and LPC abundances, most studies, including our own, have consistently provided evidence of increased Cer levels. Alterations in specific ceramide species have frequently been reported, and thus a detailed biological understanding of sphingolipid metabolism in OC warrants further efforts. Reductions in PE O- and PC O- have also been frequently reported, and thus the link between ether phospholipid metabolism and OC calls for further investigation. Our data shows the OC lipid profile can not only be distinguished from benign conditions but also from other gynecological cancers, suggesting future work focusing on identifying lipid markers unique to different gynecological malignancies may lead to lipid panels that can simultaneously test for multiple diseases. Developing clinical tests that combine lipid markers with already existing proteins markers including CA125 may provide a route for enhanced diagnosis, detection, and accurate triage of women with gynecological malignancies.

## Data Availability

Data generated in this study are available through the NIH Metabolomics Workbench (http://www.metabolomicsworkbench.org/) with project ID PR001623 (Study ID ST002521) http://dx.doi.org/10.21228/M8X42D. Code is provided at https://github.com/facundof2016/Sah_Ovarian-cancer_2023

## Acknowledgements

F.M.F and J.K. acknowledge support from NIH 1R01CA218664-01. J.K acknowledges the support from Indiana University Health–Indiana University School of Medicine Strategic Research Initiative. We acknowledge the Systems Mass Spectrometry Core at the Georgia Institute of Technology for UHPLC-MS analysis. The authors acknowledge support from NSF MRI CHE-1726528 grant for the acquisition of an ultra-high resolution Fourier transform ion cyclotron resonance (FTICR) mass spectrometer for the Georgia Institute of Technology core facilities.

## Supplementary Information

**Figure S1.**
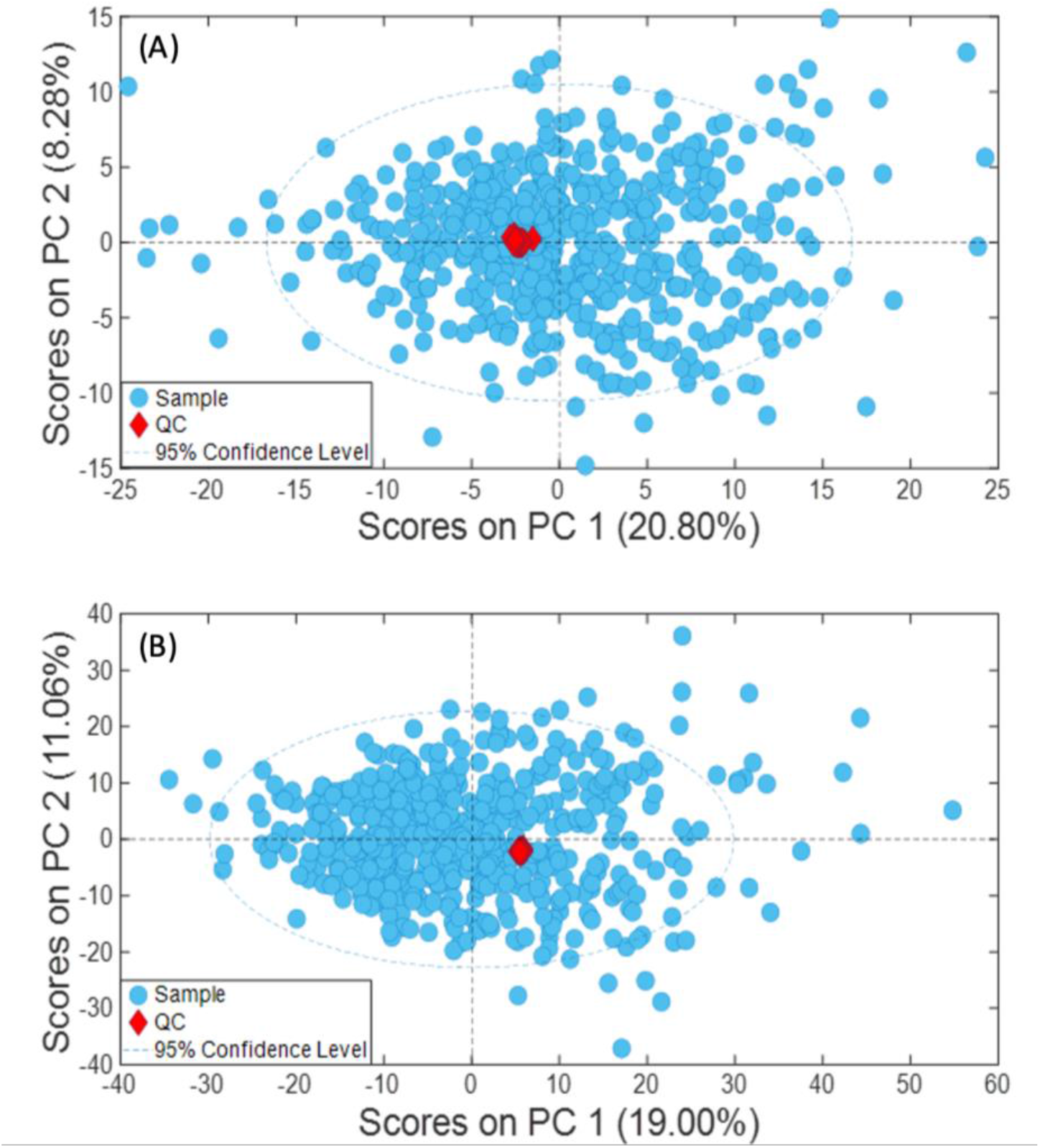
Unsupervised Principal Component Analysis (PCA) for all patient samples and quality controls. Score plot of ovarian cancer (OC) and non-OC samples, normal controls, and quality control (QC) samples using (A) 222 annotated compounds detected in (-) negative ion mode RP UHPLC-MS, (B) 780 annotated compounds detected in (+) positive ion mode RP UHPLC-MS. OC, non-OC and normal control samples are shown as blue solid circles. Quality controls are shown as red diamonds.

**Figure S2.**
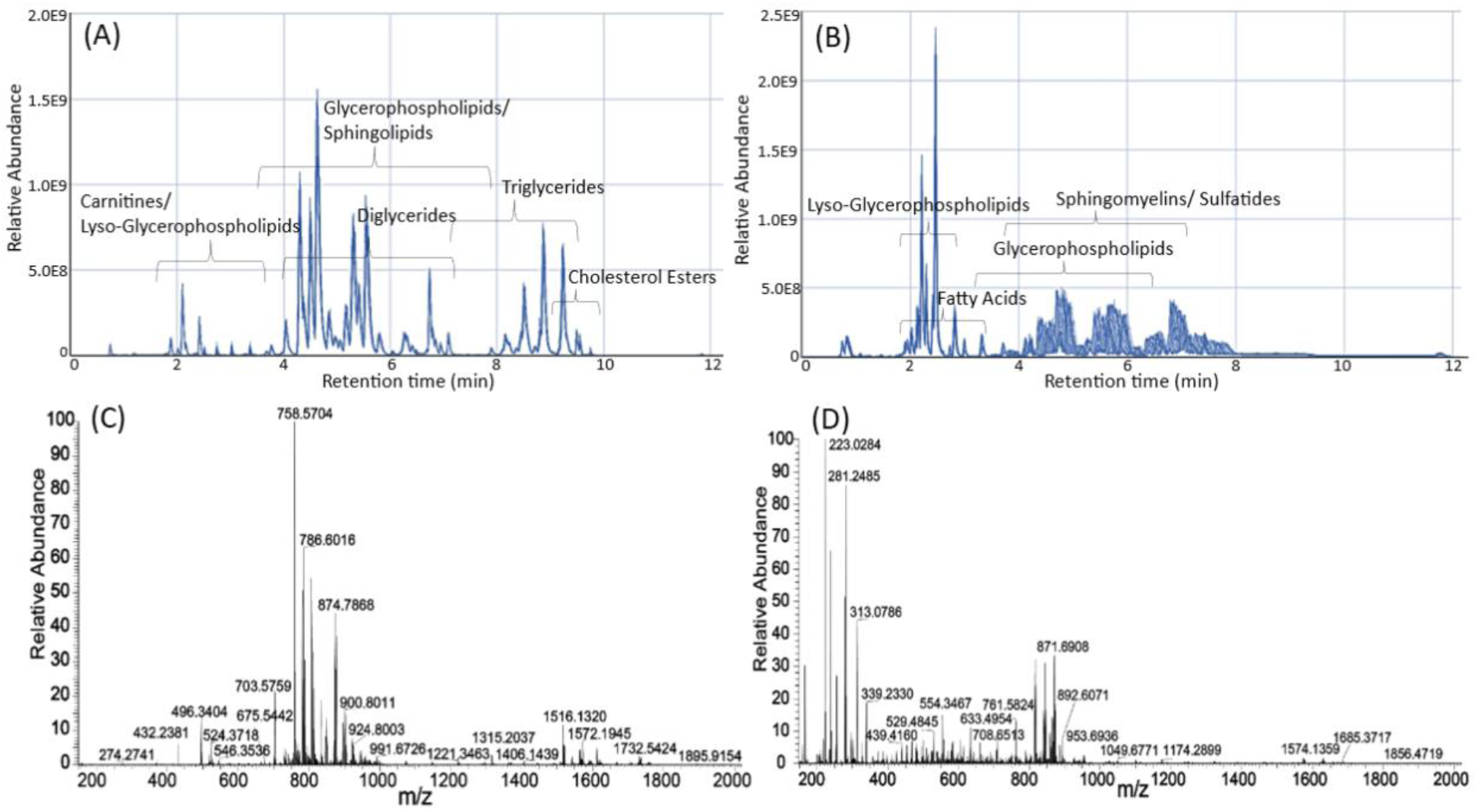
Representative chromatographic data. Overlayed total ion chromatograms (TIC) for pooled QC serum samples obtained with (a) (+) RP UHPLC-MS and (b) (-) RP UHPLC-MS. The x-axis represents the chromatographic retention time in minutes, and the y-axis peak abundance. Lipid classes are labeled based on their elution regions. Average mass spectrum for a pooled QC sample analyzed in (c) (+) RP UHPLC-MS and (b) (-) RP UHPLC-MS.

**Figure S3.**
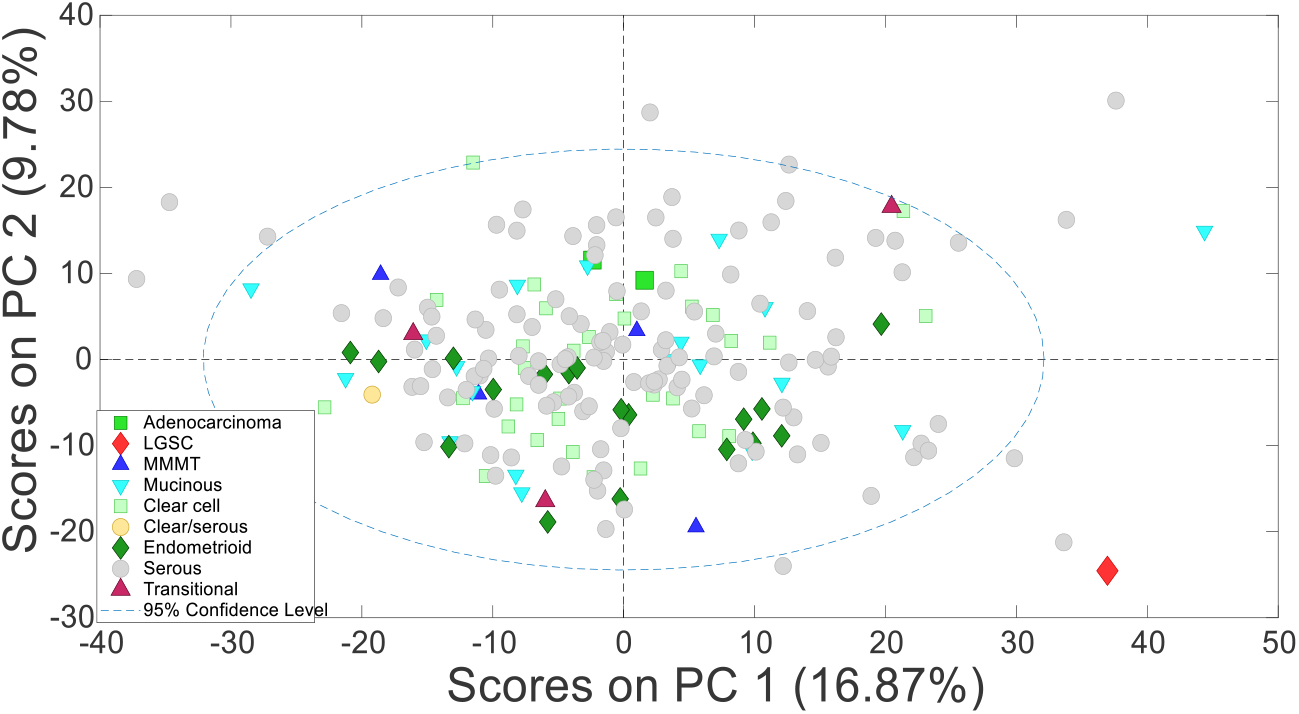
Principal Component Analysis for ovarian cancer (OC) serum samples using annotated lipid features as the input. Score plots showing clustering for OC samples of different histological types. Serous refers to high-grade serous carcinomas. LGSC : low-grade serous carcinomas, MMMT : Malignant mixed Mullerian tumor.

**Figure S4.**
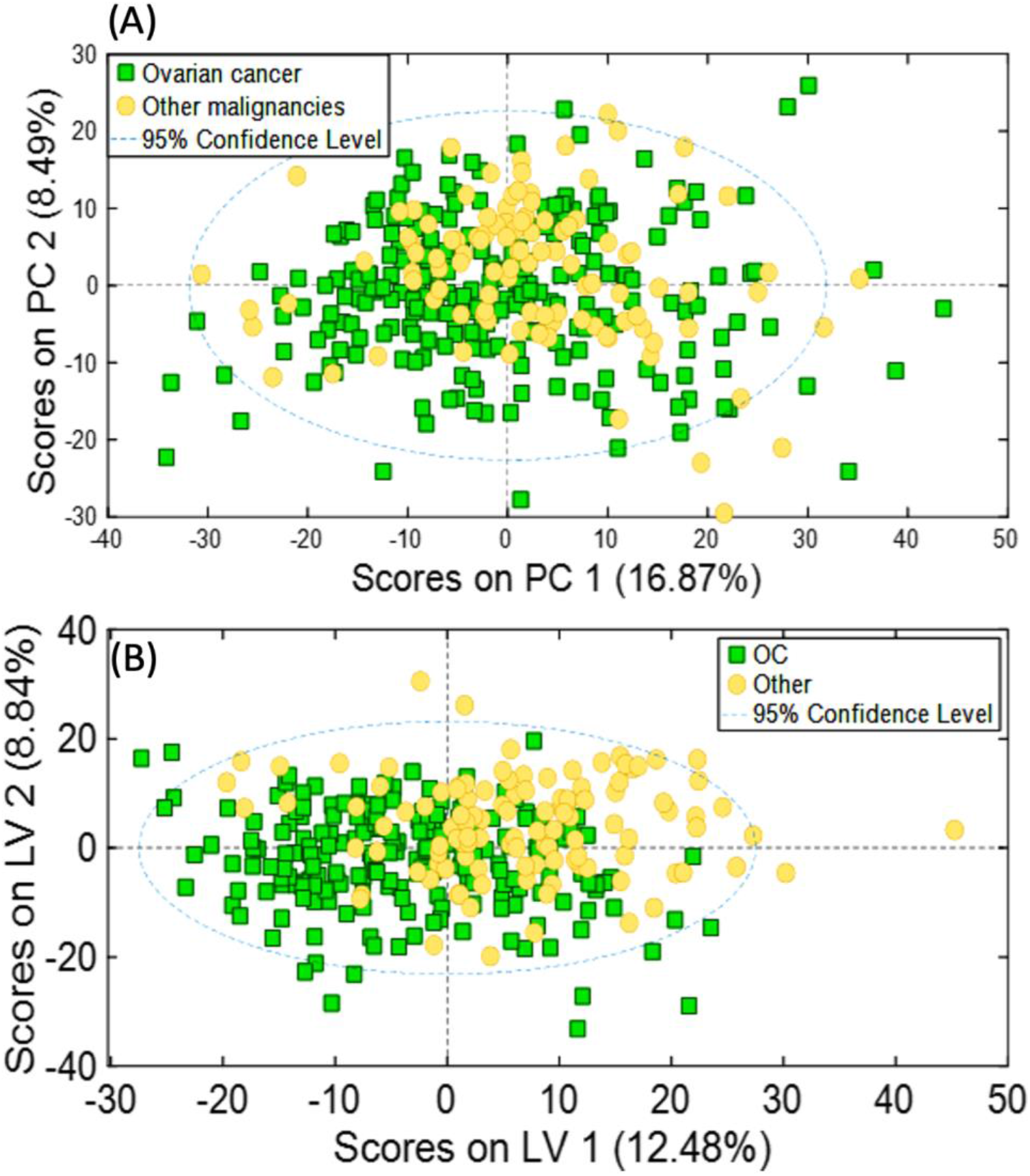
Unsupervised and supervised multivariate analysis using annotated compounds as input. (A) PCA score plot of OC patients and non-OC patients (B) oPLS-DA score plot for the same dataset OC using the combined set of (+) and (-) RP UHPLC-MS annotated features. OC samples, and non-OC samples are shown as green squares and yellow solid circles, respectively.

**Figure S5.**
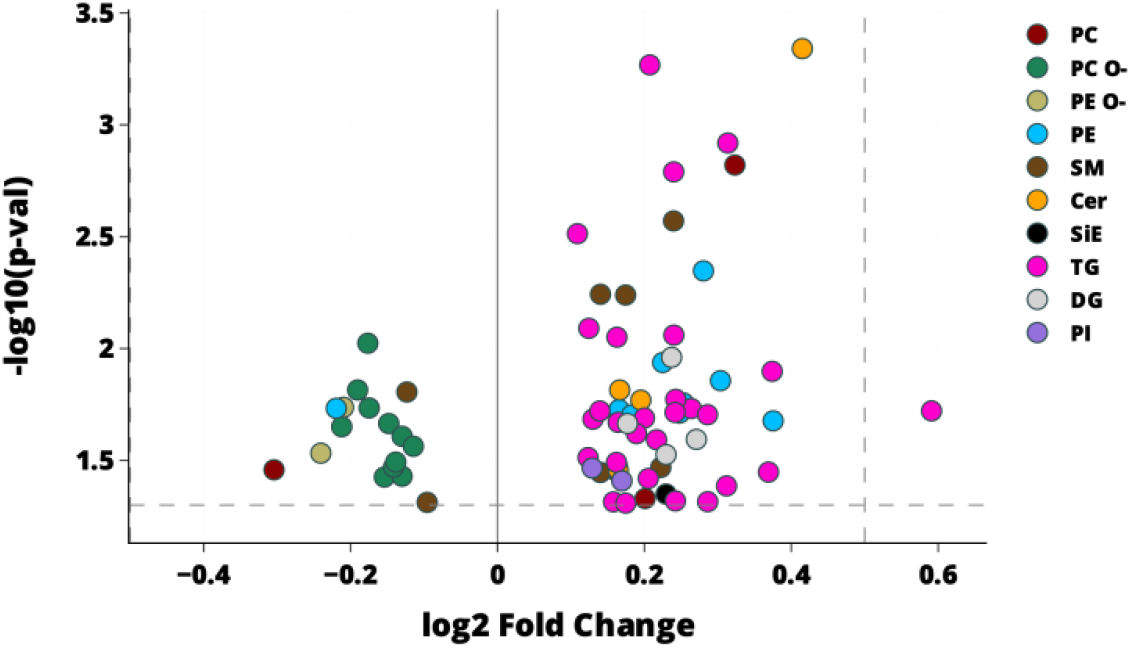
Serum lipidome analysis of high-grade serous carcinoma (HGSC) *vs.* other OC subtypes. (A) Seventy – four statisitically significant *(p*<0.05) lipids between HGSC and all other OC subtypes. The x-axis represents fold changes calculated as the base 2 logarithm of the average lipid abundance ratios for HGSC and other OC types. The y-axis shows *p*-values between HSGC and other OC types calculated as base 10 logarithm. Lipid species are color-coded by lipid class, as indicated on the plots. TG: Triacylglycerols, PC: Phosphatidylcholines, PC O-: Ether phosphatidylcholines, SM: Sphingomyelins, LPC: Lysophosphatidylcholines, Cer: Ceramides, PE O-: Ether phosphatidylethanolamines, Car: Carnitines, HexCer: Hexosylceramides, PE: Phosphatidylethanolamines, DG: Diacylglycerols, FA: Fatty acids, PI: Phosphatidylinositols, LPE: Lysophosphatidylethanolamines, PS: Phosphatidylserines, PG: Phosphatidylglycerols, MG: Monoradylglycerols, LPE O-: Ether Lysophosphatidylethanolamines, PS O- : Ether phosphatidylserines.

**Figure S6.**
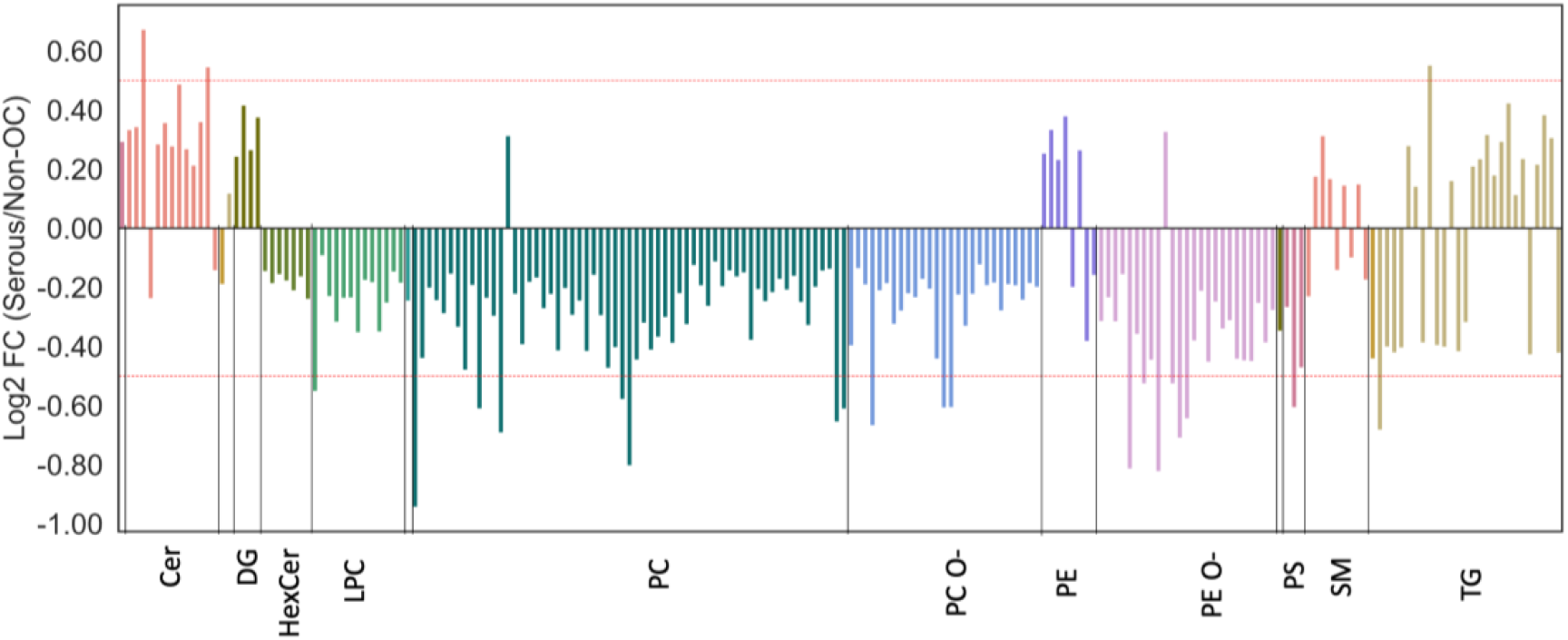
Serum lipidome analysis of HGSC vs. non-OC samples. Fold changes for statistically significant lipids with FDR corrected *p*-value < 0.05 are shown. Fold changes were calculated as the base 2 logarithm of the average abundance ratios for HGSC and non-OC conditions. A positive fold change value indicates higher levels in HGSC samples. Negative values indicate lower levels in HGSC samples. TG: Triacylglycerols, PC: Phosphatidylcholines, PC O-: Ether phosphatidylcholines, SM: Sphingomyelins, LPC: Lysophosphatidylcholines, Cer: Ceramides, PE O-: Ether phosphatidylethanolamines, HexCer: Hexosylceramides, PE: Phosphatidylethanolamines, DG: Diacylglycerols, FA: Fatty acids, PI: Phosphatidylinositols, CE: Cholesterol esters, LPE: Lysophosphatidylethanolamines, PS: Phosphatidylserines, PG: Phosphatidylglycerols, PS O- : Ether phosphatidylserines.

**Figure S7.**
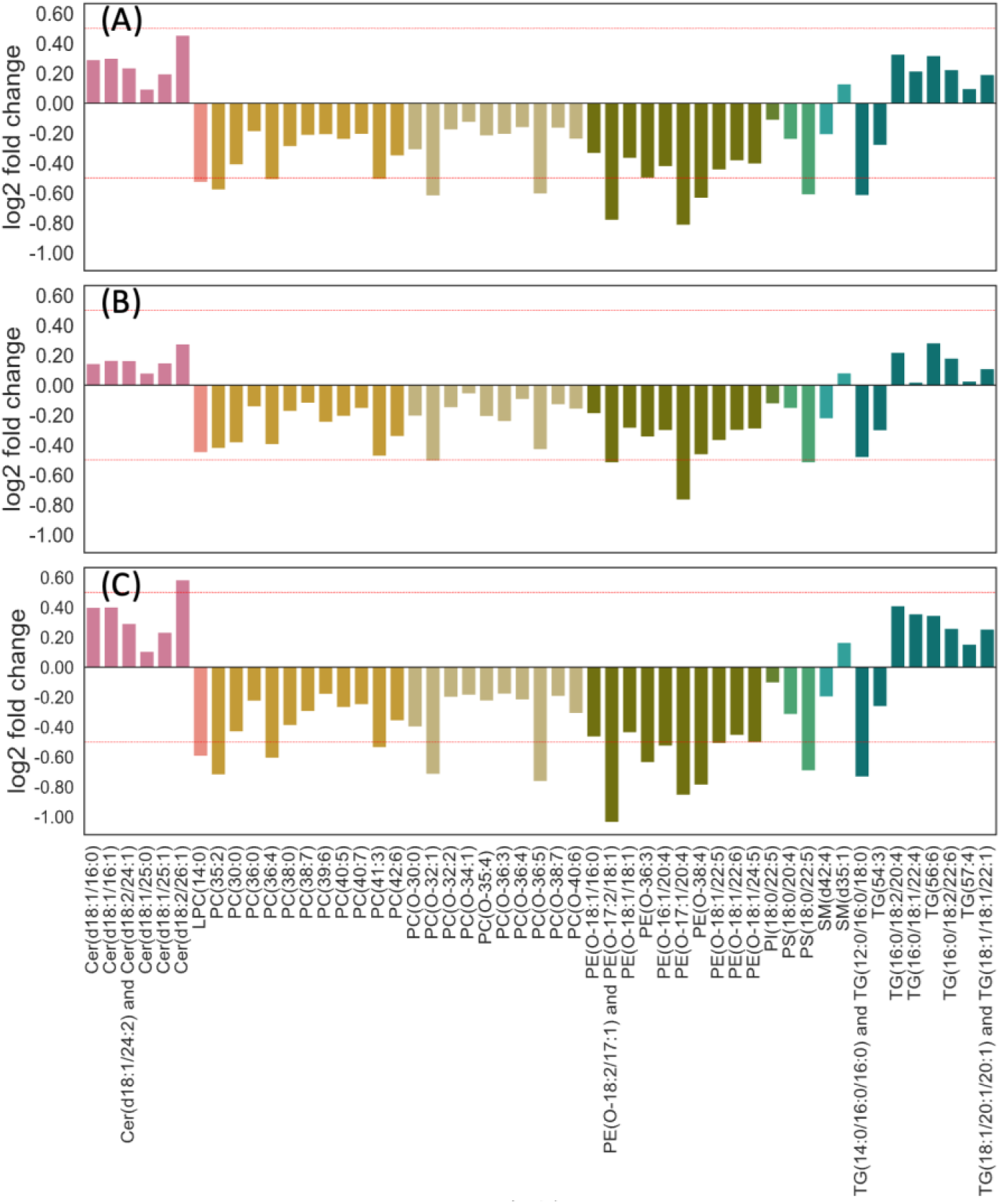
Alterations in serum lipidome using select lipid species. Fold changes for 42 lipid species selected in both early-stage *vs.* non-OC and advanced-stage *vs*. non-OC groups for (A) OC *vs*. non-OC (B) early-stage OC *vs.* non-OC and (C) advanced stage OC *vs*. non-OC. Fold changes were calculated as the base 2 logarithm of the average lipid abundance ratios for OC *vs*. non-OC. A positive fold change value indicates higher levels in OC samples. Negative values indicate lower levels in OC samples. The red dotted line indicates log2 fold change higher than 0.5 or lower than -0.5. Cer: Ceramides, LPC: Lysophosphatidylcholines, PC: Phosphatidylcholines, PC O-: Ether phosphatidylcholines, PE O-: Ether phosphatidylethanolamines, PI: Phosphatidylinositols, PS: Phosphatidylserines, SM: Sphingomyelins, TG: Triacylglycerols

**Figure S8.**
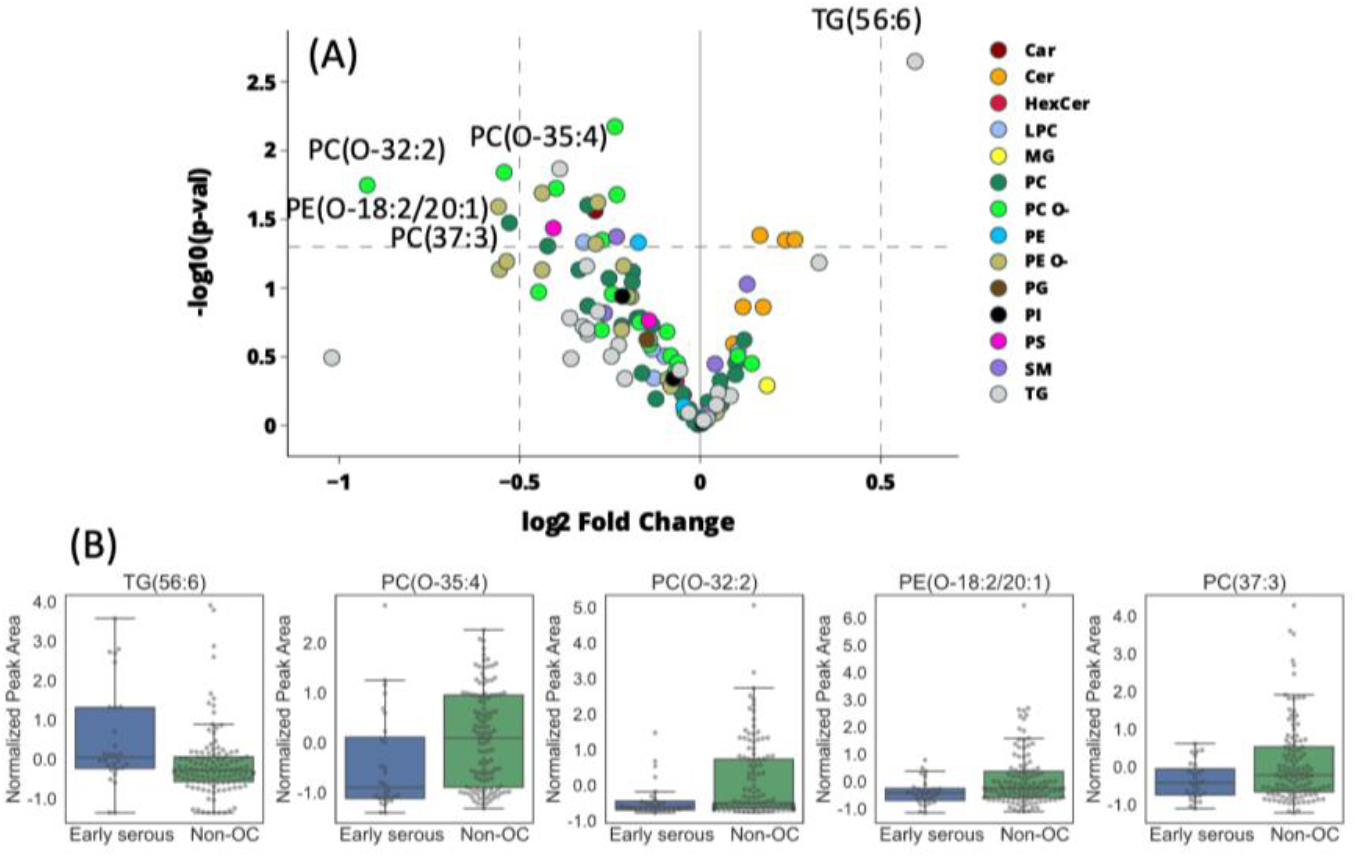
Serum lipidome alterations in early stage (I and II) HGSC. (A) Volcano plots showing lipidome differences in early-stage serous OC patients using 120 best discriminating lipids selected for early-stage OC *vs.* non-OC groups. Lipid species are color-coded by lipid class, as indicated on the plots. Fold changes were calculated as the base 2 logarithm of the average lipid abundance ratios for early serous OC *vs*. non-OC. A positive fold change value indicates higher levels in serous OC samples. Negative values indicate lower levels in serous OC samples. *p-*values were calculated using Welch’s t-test. Significantly altered lipid species (log2 fold change higher than 0.5 or lower than -0.5 and *p*-value less than 0.05) for early serous OC compared to non-OC samples are labelled. (B) Relative abundances for significantly altered lipid species. Raw data were autoscaled for visualization. TG: Triacylglycerols, PC: Phosphatidylcholines, PC O-: Ether phosphatidylcholines, SM: Sphingomyelins, LPC: Lysophosphatidylcholines, Cer: Ceramides, PE O-: Ether phosphatidylethanolamines, Car: Carnitines, HexCer: Hexosylceramides, PE: Phosphatidylethanolamines, DG: Diacylglycerols, FA: Fatty acids, PI: Phosphatidylinositols, CE: Cholesterol esters, LPE: Lysophosphatidylethanolamines, PS: Phosphatidylserines, PG: Phosphatidylglycerols, MG: Monoradylglycerols, LPE O-: Ether Lysophosphatidylethanolamines, PS O- : Ether phosphatidylserines.

**Figure S9.**
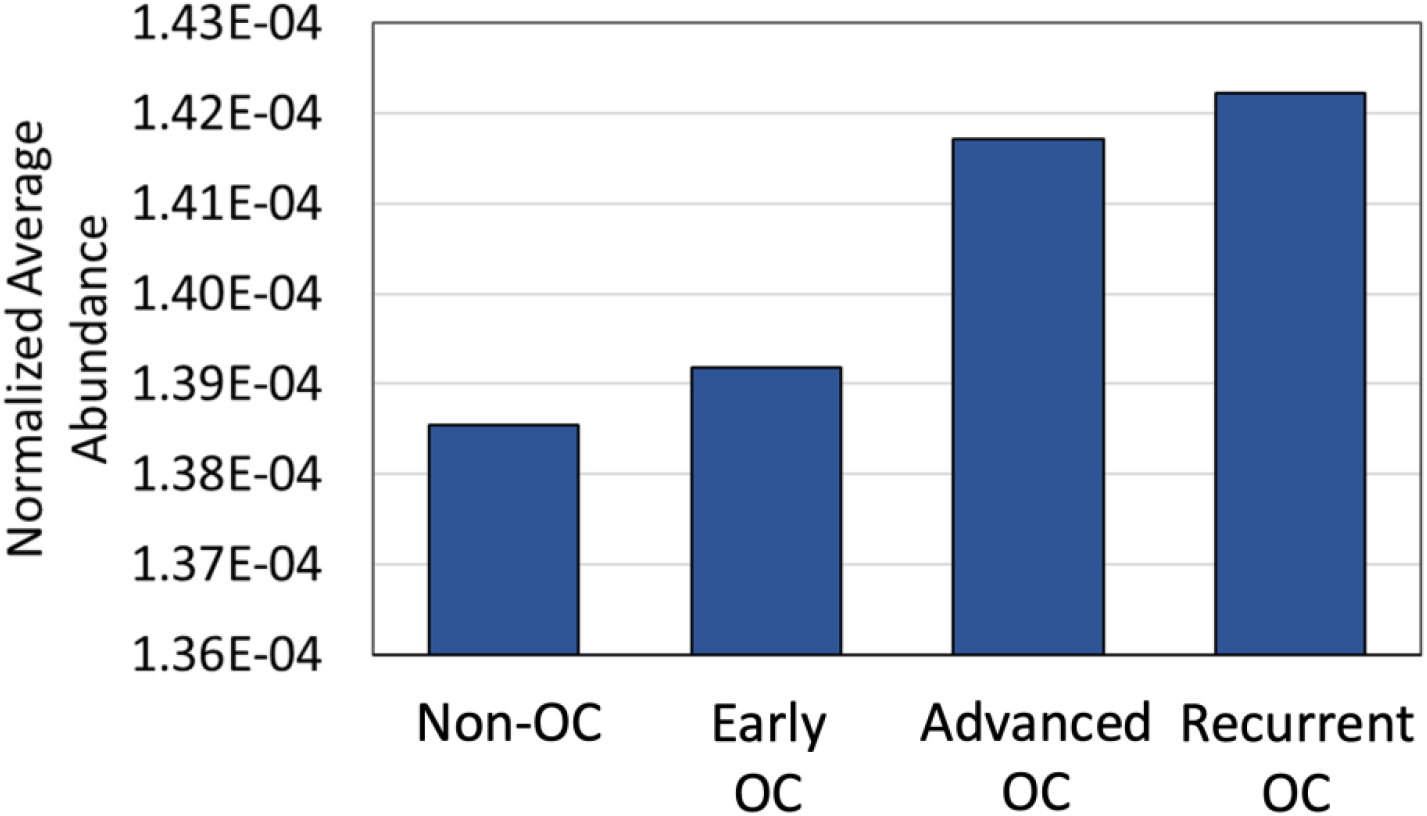
Average ganglioside abundance in OC and non-OC serum samples. For visualization purposes, data were normalized to the sum of all annotated gangliosides.

**Figure S10.**
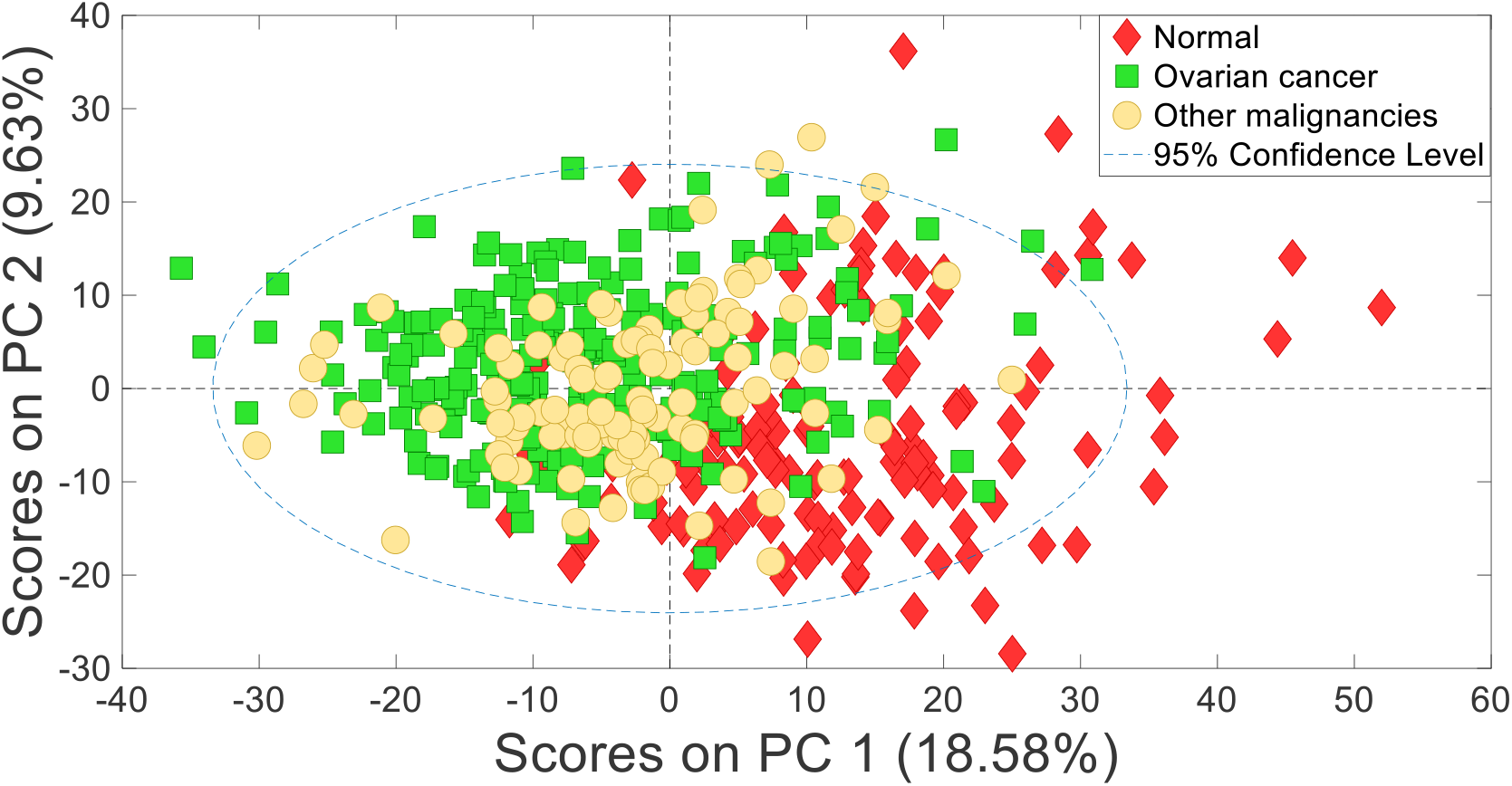
Unsupervised Principal Component Analysis (PCA) using annotated lipids as input. Score plot showing clustering of OC patients, non-OC, and normal controls using the combined set of (+) and (-) RP UHPLC-MS annotated features. OC samples, non-OC samples, normal controls are shown as green squares, yellow solid circles, and red diamonds, respectively.

**Figure S11.**
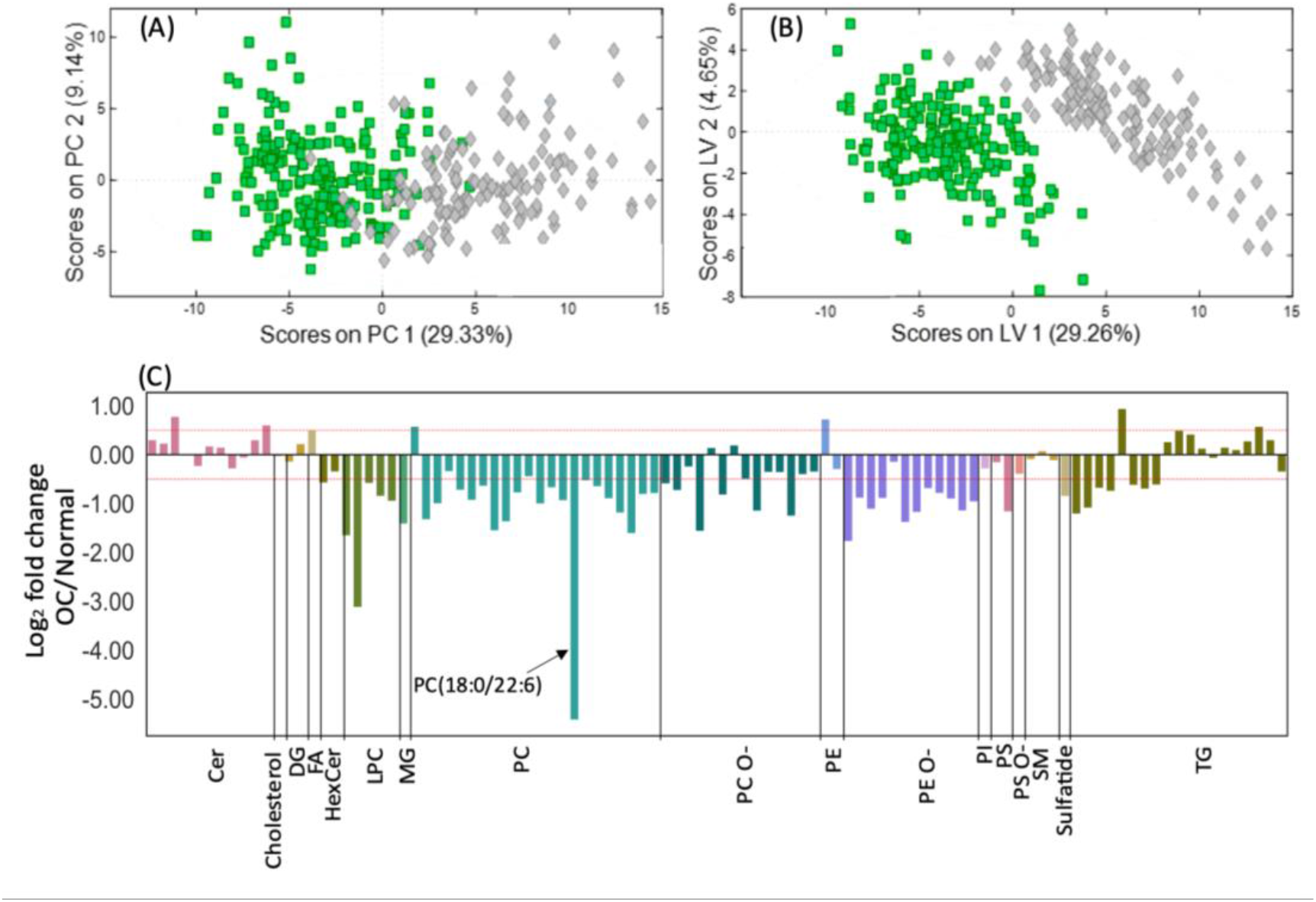
Serum lipidome analysis of OC and normal control samples using the abundances of select lipids. Lipids selected for discriminating OC and non-OC patients were used as input. One of the two highly correlated lipids was filtered out using a Pearson correlation coefficient cutoff of 0.85. Lipids with *p*-values lower than 0.05 between OC and non-OC samples were selected, followed by random forests feature selection in which lipids with a Gini index greater than the mean of all Gini indices were selected. (A) PCA score plot showing clustering of OC and normal control samples using the selected lipids. (B) o- PLS-DA score plot for the same dataset. OC samples are depicted as green squares and normal controls are shown with grey diamonds. (C) Fold changes for the selected lipids. Fold changes were calculated as the base 2 logarithm of the average lipid abundance ratios for OC *vs*. normal controls. A positive fold change value indicates higher levels in OC samples. Negative values indicate lower levels in OC samples. TG: Triacylglycerols, PC: Phosphatidylcholines, PC O-: Ether phosphatidylcholines, SM: Sphingomyelins, LPC: Lysophosphatidylcholines, Cer: Ceramides, PE O-: Ether phosphatidylethanolamines, Car: Carnitines, HexCer: Hexosylceramides, PE: Phosphatidylethanolamines, DG: Diacylglycerols, FA: Fatty acids, PI: Phosphatidylinositols, CE: Cholesterol esters, LPE: Lysophosphatidylethanolamines, PS: Phosphatidylserines, PG: Phosphatidylglycerols, MG: Monoradylglycerols, LPE O-: Ether Lysophosphatidylethanolamines, PS O- : Ether phosphatidylserines.

**Table S1.**
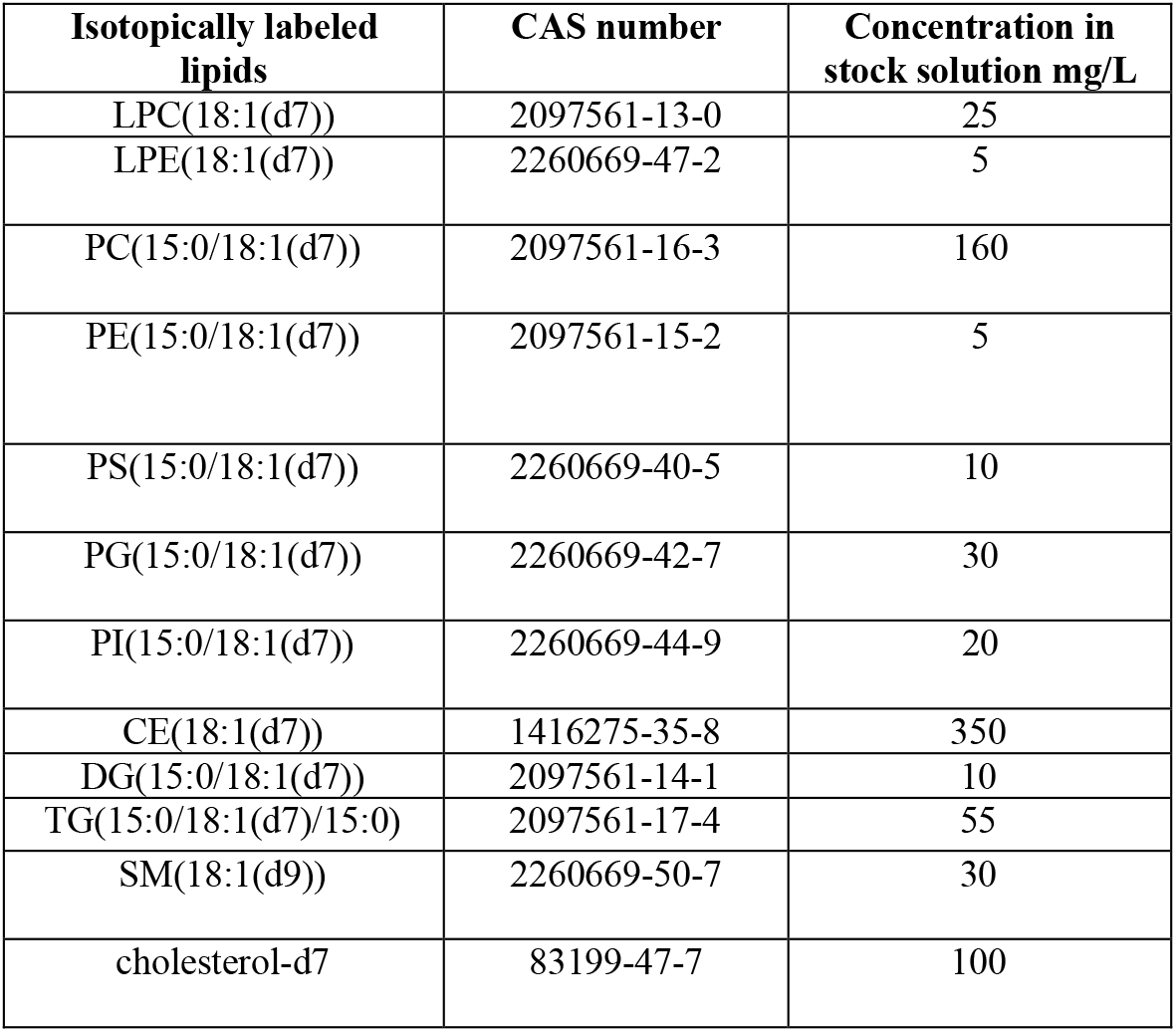
Isotopically labeled lipid internal standards (IS) used in UHPLC-MS experiments. Lipid standards for the IS mixture were purchased from Avanti Polar Lipids (Alabaster, AL).

| **Isotopically labeled lipids** | **CAS number** | **Concentration in stock solution mg/L** |
| --- | --- | --- |
| LPC(18:1(d7)) | 2097561-13-0 | 25 |
| LPE(18:1(d7)) | 2260669-47-2 | 5 |
| PC(15:0/18:1(d7)) | 2097561-16-3 | 160 |
| PE(15:0/18:1(d7)) | 2097561-15-2 | 5 |
| PS(15:0/18:1(d7)) | 2260669-40-5 | 10 |
| PG(15:0/18:1(d7)) | 2260669-42-7 | 30 |
| PI(15:0/18:1(d7)) | 2260669-44-9 | 20 |
| CE(18:1(d7)) | 1416275-35-8 | 350 |
| DG(15:0/18:1(d7)) | 2097561-14-1 | 10 |
| TG(15:0/18:1(d7)/15:0) | 2097561-17-4 | 55 |
| SM(18:1(d9)) | 2260669-50-7 | 30 |
| cholesterol-d7 | 83199-47-7 | 100 |

**Table S2.** MS parameters.

| **MS parameter** | **Positive ion mode** | **Negative ion mode** |
| --- | --- | --- |
| Capillary temperature | 275 °C | 275 °C |
| Spray voltage | +3.0 kV | -2.8 kV |
| Sheath gas flow rate | 60 Arb. | 60 Arb. |
| Auxiliary gas flow rate | 18 Arb. | 18 Arb. |
| Sweep gas flow rates | 4 Arb. | 4 Arb. |
| Probe Heater Temperature | 425 °C | 425 °C |

Arb: Arbitrary units.

**Table S3.** Chromatographic gradient used for UHPLC-MS. For negative ion mode UHPLC-MS mobile phase A was 10 mM ammonium acetate with water/acetonitrile (40:60 v/v) and mobile phase B was 10 mM ammonium acetate with 2-isopropanol/acetonitrile (90:10 v/v). For positive ion mode, mobile phase A was 10 mM ammonium formate with water/acetonitrile (40:60 v/v) and 0.1% formic acid. Mobile phase B was 10 mM ammonium formate with 2-isopropanol/acetonitrile (90:10 v/v) and 0.1% formic acid.

| **RP UHPLC Gradient** |  |  |  |
| --- | --- | --- | --- |
| **Time (min)** | **Mobile phase A** | **Mobile phase B** | **Flow rate (ml min<sup>-1</sup>)** |
| 0.0 | 80% | 20% | 0.4 |
| 0.0 | 80% | 20% | 0.4 |
| 1.0 | 40% | 60% | 0.4 |
| 5.0 | 30% | 70% | 0.4 |
| 5.5 | 15% | 85% | 0.4 |
| 8.0 | 10% | 90% | 0.4 |
| 8.2 | 0% | 100% | 0.4 |
| 10.5 | 0% | 100% | 0.4 |
| 10.7 | 80% | 20% | 0.4 |
| 12.0 | 80% | 20% | 0.4 |

**Table S4.** UHPLC-MS system suitability. Percent relative standard deviation (%RSD) of the internal standards (IS) spiked in pooled QC injections in (a) (-) RP UHPLC-MS and (b) (+) RP UHPLC-MS experiments. %RSD were calculated in Compound Discoverer. In cases where Compound Discoverer failed to detect some of the IS, data were evaluated manually using Xcalibur (Thermo Scientific).

| (A) |  |  |  |  |  |  |  |  |
| --- | --- | --- | --- | --- | --- | --- | --- | --- |
| Isotopically Labeled IS | Adduct | Experimental Exact Mass | Theoretical Exact Mass | Mass Error (ppm) | Retention Time (min) | Processed in Compound Discoverer | Detected in Xcalibur | Compound Discoverer %RSD QC (n=54) |
| LPC(18:1(d7)/0:0) | [M+HCOOH-H]<sup>-</sup> | 574.3978 | 574.3965 | 0.40 | 2.19 | Yes | Yes | 6 |
| PC(33:1(d7))>PC(15:0/18:1(d7)) | [M+HCOOH-H]<sup>-</sup> | 798.6119 | 798.6122 | 0.40 | 5.10 | Yes | Yes | 2 |
| LPE(18:1(d7)) | [M-H]<sup>-</sup> | 486.3454 | 486.3442 | 0.57 | 2.21 | Yes | Yes | 2 |
| PE(15:0/18:1(d7)) | [M-H]<sup>-</sup> | 710.5593 | 710.5588 | 0.27 | 5.08 | Yes | Yes | 7 |
| PG(15:0/18:1(d7)) | [M-H]<sup>-</sup> | 741.5539 | 741.5539 | 0.19 | 3.92 | Yes | Yes | 4 |
| PI(15:0/18:1(d7)) | [M-H]<sup>-</sup> | 829.5696 | 829.5696 | -0.19 | 3.80 | Yes | Yes | 2 |
| PS(15:0/18:1(d7)) | [M-H]<sup>-</sup> | 754.5492 | 754.5490 | 0.25 | 3.93 | Yes | Yes | 1 |
| SM(d18:1/18:1(d9)) | [M+HCOOH-H]<sup>-</sup> | 783.6452 | 783.6450 | 0.20 | 4.53 | No | Yes | NA |
| (B) |  |  |  |  |  |  |  |  |
| Isotopically Labeled IS | Adduct | Experimental Exact Mass | Theoretical Exact Mass | Mass Error (ppm) | Retention Time (min) | Processed in Compound Discoverer | Detected in Xcalibur | Compound Discoverer %RSD QC (n=54) |
| DG(15:0/18:1(d7)/0:0) | [M+NH<sub>4</sub>]<sup>+</sup> | 604.5768 | 604.5700 | 0.36 | 6.49 | Yes | Yes | 3 |
| LPC(18:1(d7)/0:0) | [M+H]<sup>+</sup> | 528.3921 | 528.3921 | 0.01 | 2.16 | Yes | Yes | 1 |
| PC(15:0/18:1(d7)) | [M+H]<sup>+</sup> | 752.6068 | 752.6061 | 0.95 | 4.84 | Yes | Yes | 3 |
| LPE(18:1(d7)) | [M+H]<sup>+</sup> | 486.3454 | 486.3451 | 0.54 | 2.19 | Yes | Yes | 3 |
| PE(15:0/18:1(d7)) | [M+H]<sup>+</sup> | 710.5599 | 710.5591 | 1.05 | 5.03 | Yes | Yes | 1 |
| PI(15:0/18:1(d7)) | [M+NH<sub>4</sub>]<sup>+</sup> | 846.5967 | 846.5960 | 0.42 | 4.65 | Yes | Yes | 15 |
| PS(15:0/18:1(9z,d7)) | [M+H]<sup>+</sup> | 754.5494 | 754.5490 | 0.53 | 4.27 | Yes | Yes | 19 |
| SM(d18:1/18:1(d9)) | [M+H]<sup>+</sup> | 737.6400 | 737.6397 | 0.42 | 4.44 | Yes | Yes | 1 |
| TG(15:0/18:1(d7)/15:0) | [M+NH<sub>4</sub>]<sup>+</sup> | 828.7916 | 828.7915 | 0.11 | 8.81 | Yes | Yes | 1 |
| CE(18:1(d7)) | [M+NH<sub>4</sub>]<sup>+</sup> | 674.6706 | 674.6700 | 0.85 | 9.75 | No | Yes | NA |
| Cholesterol-d7 | [M+H<sub>2</sub>O+H]<sup>+</sup> | NA | 375.3800 | NA | 4.39 | No | No | NA |
| PG(15:0/18:1(d7)) | [M+H]<sup>+</sup> | 741.5537 | 741.5525 | 1.58 | 4.10 | No | Yes | NA |

**Table S5:** Annotated gangliosides. Proposed annotation, experimental monoisotopic *m/z* value, mass error, retention time, and MS annotation confidence level are provided. MS annotation level was assigned based on the following criteria: 1) MS1 and MS/MS spectrum of standard matched to the feature. 2) MS1 and MS/MS spectrum of the feature matched with library spectra 3) tentative ID assignment based on elemental formula match with literature. 4) unknowns

| Name | Adduct | Experimental $m/z$ | Mass error ppm | Retention time min | Annotation level |
| --- | --- | --- | --- | --- | --- |
| asialo-GM1(d18:1/24:0) | [M+H]<sup>+</sup> | 1339.8835 | 0.76 | 6.83 | 2 |
| asialo-GM1(d18:1/16:0) | [M+H]<sup>+</sup> | 1227.7578 | 0.40 | 3.97 | 2 |
| GM3(d18:1/18:0) | [M-H]<sup>-</sup> | 1179.7358 | 0.77 | 3.95 | 2 |
| GM3(d18:1/16:0) | [M-H]<sup>-</sup> | 1151.7043 | -1.32 | 3.36 | 2 |
| GM3(d18:1/18:1) | [M-H]<sup>-</sup> | 1177.7201 | -1.13 | 3.44 | 2 |
| GM3(d40:1) | [M-H]<sup>-</sup> | 1235.7988 | -0.78 | 5.57 | 2 |
| GM3(d42:1) | [M-H]<sup>-</sup> | 1263.8300 | -0.74 | 6.39 | 2 |
| GM1(d18:1/16:0) | [M+H]<sup>+</sup> | 1518.8530 | 0.26 | 3.64 | 2 |
| GM3(d18:0/24:2) | [M+H]<sup>+</sup> | 1263.8311 | 0.24 | 6.33 | 2 |
| GM3(d18:1/12:0) | [M+H]<sup>+</sup> | 1097.6578 | -0.009 | 2.80 | 3 |
| GM3(d18:1/14:0) | [M+H]<sup>+</sup> | 1125.6891 | 0.053 | 3.18 | 2 |
| GM3(d18:1/16:0) | [M+H]<sup>+</sup> | 1153.7204 | 0.00 | 3.70 | 2 |
| GM3(d18:1/16:1) | [M+H]<sup>+</sup> | 1151.7048 | 0.26 | 3.24 | 2 |
| GM3(d18:1/18:0) | [M+H]<sup>+</sup> | 1181.7526 | 0.76 | 4.40 | 2 |
| GM3(d18:1/18:1) | [M+H]<sup>+</sup> | 1179.7362 | 0.12 | 3.81 | 3 |
| GM3(d18:1/20:0) | [M+H]<sup>+</sup> | 1209.7836 | 0.50 | 5.26 | 3 |
| GM3(d18:1/22:0) | [M+H]<sup>+</sup> | 1237.8147 | 0.32 | 6.17 | 2 |
| GM3(d18:1/22:1) | [M+H]<sup>+</sup> | 1235.7997 | 0.83 | 5.19 | 3 |
| GM3(d18:1/22:1) | [M+H]<sup>+</sup> | 1235.8000 | 1.11 | 5.43 | 3 |
| GM3(d18:1/24:0) | [M+H]<sup>+</sup> | 1265.8463 | 0.55 | 6.70 | 2 |
| GM3(d18:1/24:1) | [M+H]<sup>+</sup> | 1263.8303 | 0.25 | 6.05 | 2 |
| GM3(d18:1/24:2) | [M+H]<sup>+</sup> | 1261.8155 | 1.00 | 5.31 | 3 |

**Table S6.** Best differential lipids selected for advanced-stage OC (III and IV) *vs.* non-OC. *p*-values were calculated with Welch’s t-test. Fold changes were calculated as the base 2 logarithm of the average lipid abundance ratios for advanced OC *vs*. non-OC samples. A positive fold change value indicates higher levels in OC samples. Negative values indicate lower levels in OC samples. TG: Triacylglycerols, PC: Phosphatidylcholines, PC O-: Ether phosphatidylcholines, SM: Sphingomyelins, LPC: Lysophosphatidylcholines, Cer: Ceramides, PE O-: Ether phosphatidylethanolamines, Car: Carnitines, HexCer: Hexosylceramides, PE: Phosphatidylethanolamines, DG: Diacylglycerols, FA: Fatty acids, PI: Phosphatidylinositols, CE: Cholesterol esters, LPE: Lysophosphatidylethanolamines, PS: Phosphatidylserines, PG: Phosphatidylglycerols, MG: Monoradylglycerols, LPE O-: Ether Lysophosphatidylethanolamines, PS O- : Ether phosphatidylserines.

| Lipid class | Annotation | T-test p-value | Log2 fold change (OC/non-OC) | Retention Time (minutes) |
| --- | --- | --- | --- | --- |
| Car | Car(18:1) | 3.41E-03 | 0.16 | 2.10 |
| Cer | Cer(d18:1_16:1) | 1.00E-04 | 0.40 | 5.32 |
| Cer | Cer(d18:1_16:0) | 2.92E-11 | 0.40 | 5.34 |
| Cer | Cer(d18:0_18:1) | 6.17E-12 | 0.77 | 6.31 |
| Cer | Cer(d18:1_24:2) and Cer(d18:2_24:1) | 2.52E-07 | 0.29 | 6.83 |
| Cer | Cer(d18:1_24:2) and Cer(d18:2_24:1) | 2.38E-06 | 0.53 | 6.93 |
| Cer | Cer(d18:2_23:0) | 3.37E-07 | 0.30 | 6.94 |
| Cer | Cer(d18:2_26:1) | 7.12E-10 | 0.58 | 7.12 |
| Cer | Cer(d18:0_22:0) | 4.59E-05 | -0.32 | 7.14 |
| Cer | Cer(d18:1_25:1) | 1.68E-04 | 0.27 | 7.16 |
| Cer | Cer(d18:1_25:1) | 1.50E-03 | 0.23 | 7.23 |
| Cer | Cer(d18:1_26:1) | 1.08E-08 | 0.41 | 7.38 |
| Cer | Cer(d18:1_25:0) | 4.76E-02 | 0.11 | 7.63 |
| Cholesterol | Cholesterol | 1.37E-04 | 0.13 | 4.99 |
| Cholesterol sulfate | Cholesterol sulfate | 5.26E-05 | -0.21 | 2.59 |
| DG | DG(18:2_18:2_0:0) | 8.44E-04 | 0.32 | 5.62 |
| DG | DG(18:0_20:4_0:0) | 2.54E-03 | 0.24 | 5.63 |
| DG | DG(35:4) | 3.97E-05 | 0.30 | 6.08 |
| FA | FA(22:1) | 8.18E-03 | 0.19 | 3.35 |
| HexCer | HexCer(d18:1_22:0) | 6.36E-03 | -0.15 | 6.84 |
| LPC | LPC(14:0) | 5.14E-08 | -0.60 | 1.64 |
| LPC | LPC(17:1) | 1.58E-03 | -0.25 | 1.99 |
| LPC | LPC(22:0) | 1.12E-07 | -0.41 | 3.28 |
| PC | PC(41:3) | 1.65E-02 | -0.54 | 3.52 |
| PC | PC(28:0) | 3.04E-04 | -0.98 | 3.71 |
| PC | PC(41:7) | 2.79E-05 | -0.46 | 3.79 |
| PC | PC(40:8) | 1.02E-11 | -0.51 | 3.81 |
| PC | PC(38:7) | 3.26E-05 | -0.30 | 3.95 |
| PC | PC(36:4) | 1.24E-04 | -0.61 | 4.11 |
| PC | PC(39:6) | 1.63E-02 | -0.18 | 4.29 |
| PC | PC(30:0) | 4.66E-07 | -0.43 | 4.39 |
| PC | PC(40:7) | 1.99E-02 | -0.25 | 4.47 |
| PC | PC(32:1) | 4.49E-02 | -0.15 | 4.47 |
| PC | PC(39:6) | 9.67E-03 | -0.31 | 4.57 |
| PC | PC(42:8) | 1.33E-02 | -0.12 | 4.96 |
| PC | PC(42:6) | 1.35E-03 | -0.44 | 5.15 |
| PC | PC(40:5) | 1.92E-03 | -0.27 | 5.44 |
| PC | PC(35:0) | 3.87E-04 | -0.39 | 5.66 |
| PC | PC (35:2) | 1.07E-09 | -0.72 | 5.72 |
| PC | PC(39:4) | 3.58E-04 | -0.22 | 5.89 |
| PC | PC(42:6) | 1.12E-03 | -0.36 | 6.16 |
| PC | PC(35:2) | 6.84E-03 | -0.35 | 6.49 |
| PC | PC(36:0) | 1.06E-04 | -0.23 | 6.76 |
| PC | PC(38:0) | 6.76E-03 | -0.39 | 7.08 |
| PC | PC(16:0_22:5) and PC(18:1_20:4) | 2.68E-03 | -0.15 | 4.70 |
| PC | PC(38:6) | 1.68E-05 | -0.27 | 4.12 |
| PC | PC(18:0_22:6) | 7.75E-05 | -0.19 | 4.79 |
| PC O- | PC(O-35:4) | 4.55E-02 | -0.23 | 4.61 |
| PC O- | PC(O-38:7) | 5.36E-04 | -0.22 | 4.66 |
| PC O- | PC(O-30:0) | 8.20E-10 | -0.40 | 4.97 |
| PC O- | PC(O-32:2) | 1.44E-04 | -0.20 | 5.00 |
| PC O- | PC(O-38:7) | 4.88E-06 | -0.20 | 5.03 |
| PC O- | PC(O-36:4) | 8.73E-05 | -0.22 | 5.05 |
| PC O- | PC(O-32:1) | 7.42E-06 | -0.72 | 5.32 |
| PC O- | PC(O-36:3) | 5.81E-02 | -0.18 | 5.65 |
| PC O- | PC(O-40:6) | 1.99E-07 | -0.31 | 5.77 |
| PC O- | PC(O-38:3) | 1.63E-04 | -0.35 | 6.40 |
| PC O- | PC(O-36:5) | 1.82E-08 | -0.77 | 6.49 |
| PC O- | PC(O-34:1) | 5.99E-04 | -0.19 | 6.59 |
| PC O- | PC(O-16:0_18:2) | 9.02E-08 | -0.39 | 5.55 |
| PC O- | PC(O-36:3) | 6.38E-08 | -0.28 | 5.79 |
| PC O- | PC(O-38:6) | 7.57E-08 | -0.35 | 5.07 |
| PC O- | PC(O-20:7_18:0) and PC(O-16:1_22:6) | 1.89E-05 | -0.23 | 4.86 |
| PC O- | PC(O-18:1_22:6) and PC(O-22:7_18:0) | 4.60E-09 | -0.71 | 5.85 |
| PC P- | PC(P-18:1_20:4) | 1.95E-03 | -0.22 | 6.17 |
| PE | PE(16:0_22:6) | 2.63E-07 | 0.34 | 3.10 |
| PE O- | PE(O-17:1_20:4) | 5.85E-06 | -0.86 | 3.79 |
| PE O- | PE(O-17:1_20:4) | 1.26E-05 | 0.40 | 4.09 |
| PE O- | PE(O-16:1_20:4) | 1.85E-10 | -0.53 | 5.13 |
| PE O- | PE(O-36:3) | 2.59E-09 | -0.64 | 5.32 |
| PE O- | PE(O-18:2_17:1) and PE(O-17:2_18:1) | 3.87E-10 | -1.04 | 5.79 |
| PE O- | PE(O-18:1_22:6) | 3.56E-09 | -0.46 | 5.83 |
| PE O- | PE(O-18:1_22:5) | 1.31E-07 | -0.54 | 6.06 |
| PE O- | PE(O-38:4) | 8.46E-11 | -0.79 | 6.26 |
| PE O- | PE(O-18:1_22:5) | 7.08E-08 | -0.51 | 6.41 |
| PE O- | PE(O-18:1_16:0) | 5.05E-07 | -0.47 | 6.68 |
| PE O- | PE(O-18:1_18:1) | 3.04E-08 | -0.44 | 6.70 |
| PE O- | PE(O-18:1_22:4) | 3.59E-07 | -0.36 | 6.72 |
| PE O- | PE(O-18:1_24:5) | 7.69E-05 | -0.51 | 6.84 |
| PE O- | PE(O-18:1_22:4) | 1.69E-07 | -0.38 | 6.99 |
| PE O- | PE(O-18:0_20:4) and PE(O-18:2_20:2) | 2.75E-03 | -0.18 | 5.35 |
| PI | PI(18:0_22:5) | 9.40E-02 | -0.11 | 5.04 |
| PS | PS(18:0_20:4) | 2.67E-07 | -0.32 | 4.80 |
| PS | PS(18:0_22:5) | 4.33E-08 | -0.69 | 3.76 |
| PS O- | PS(O-20:3_20:4) | 6.63E-05 | -0.46 | 5.24 |
| SM | SM(d40:3) | 2.44E-03 | 0.15 | 5.38 |
| SM | SM(d42:4) | 5.94E-03 | -0.20 | 6.81 |
| SM | SM(d35:1) | 1.57E-03 | 0.17 | 5.03 |
| TG | TG(16:0_18:2_22:6) | 2.93E-02 | 0.26 | 8.10 |
| TG | TG(18:2_18:2_22:4) | 5.27E-04 | 0.31 | 8.35 |
| TG | TG(16:0_18:2_20:4) | 7.28E-05 | 0.41 | 8.41 |
| TG | TG(14:0_16:0_18:2) and TG(14:0_16:1_18:1) | 2.41E-03 | -0.39 | 8.45 |
| TG | TG(18:0/18:1/18:2) and TG(18:1/18:1/18:1) and TG(16:0/18:2/20:1) | 5.46E-03 | -0.26 | 8.47 |
| TG | TG(57:4) | 5.66E-03 | 0.15 | 8.51 |
| TG | TG(18:0_20:4_20:4) and TG(18:0_18:2_22:6) | 1.82E-04 | 0.45 | 8.52 |
| TG | TG(18:1_18:1_20:4) and TG(16:0_18:1_22:5) and TG(18:1_18:2_20:3) | 1.41E-03 | 0.26 | 8.72 |
| TG | TG(18:1_18:1_20:4) and TG(16:0_18:1_22:5) and TG(18:1_18:2_20:3) | 1.32E-02 | 0.35 | 8.78 |
| TG | TG(16:0_17:0_18:0) and TG(16:0_16:0_19:0) and TG(15:0_17:0_19:0) | 1.87E-04 | 0.20 | 8.87 |
| TG | TG(18:1_20:1_20:1) and TG(18:1_18:1_22:1) | 8.40E-07 | 0.26 | 8.87 |
| TG | TG(14:0_16:0_16:0) and TG(12:0_16:0_18:0) | 4.32E-05 | -0.73 | 8.87 |
| TG | TG(16:0_18:1_19:0) and TG(17:0_18:0_18:1) | 1.73E-02 | -0.32 | 8.90 |
| TG | TG(16:0_18:1_22:4) | 3.66E-06 | 0.36 | 9.16 |
| TG | TG(60:4) | 1.19E-01 | -0.24 | 9.64 |

**Table S7.**
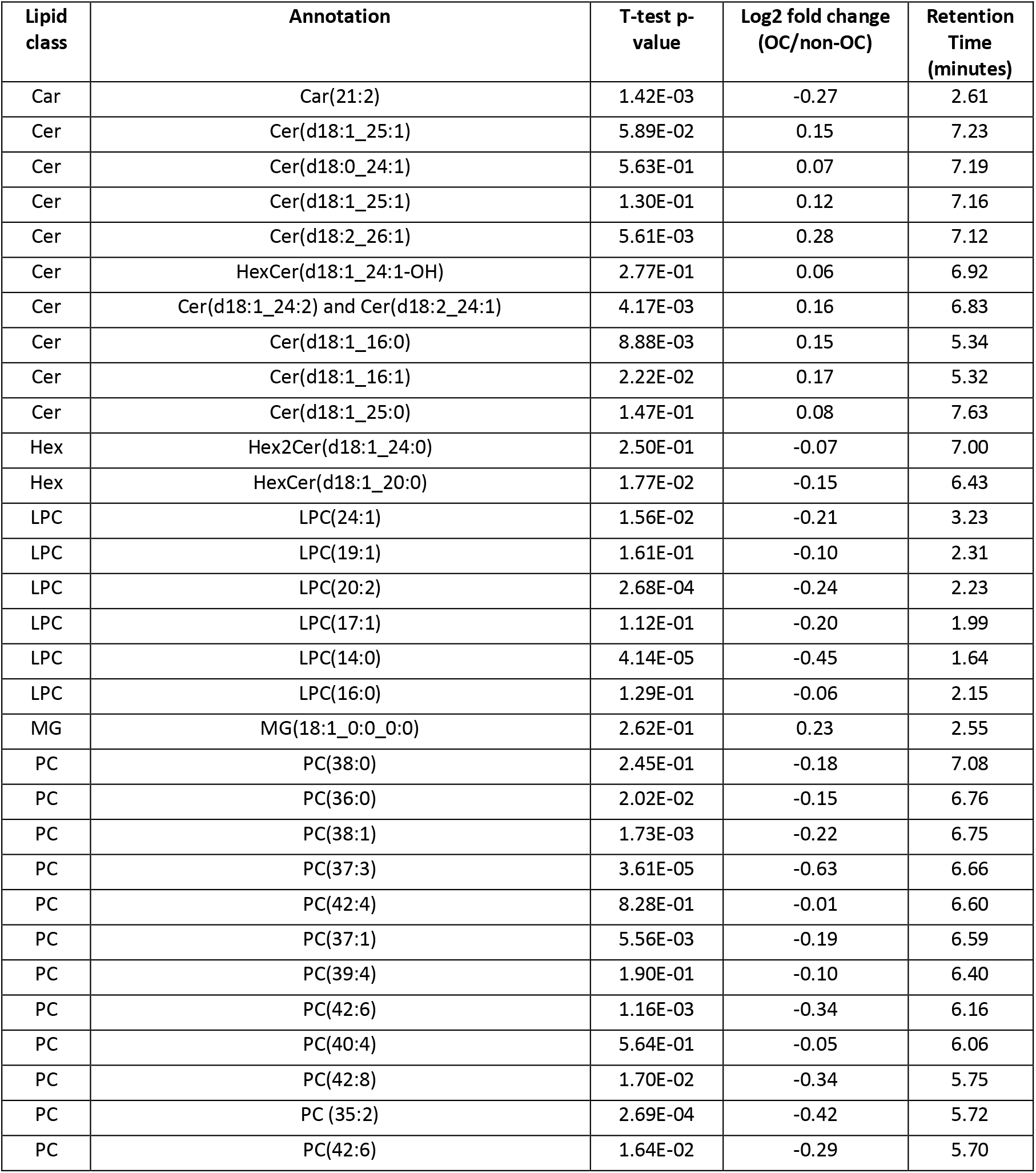
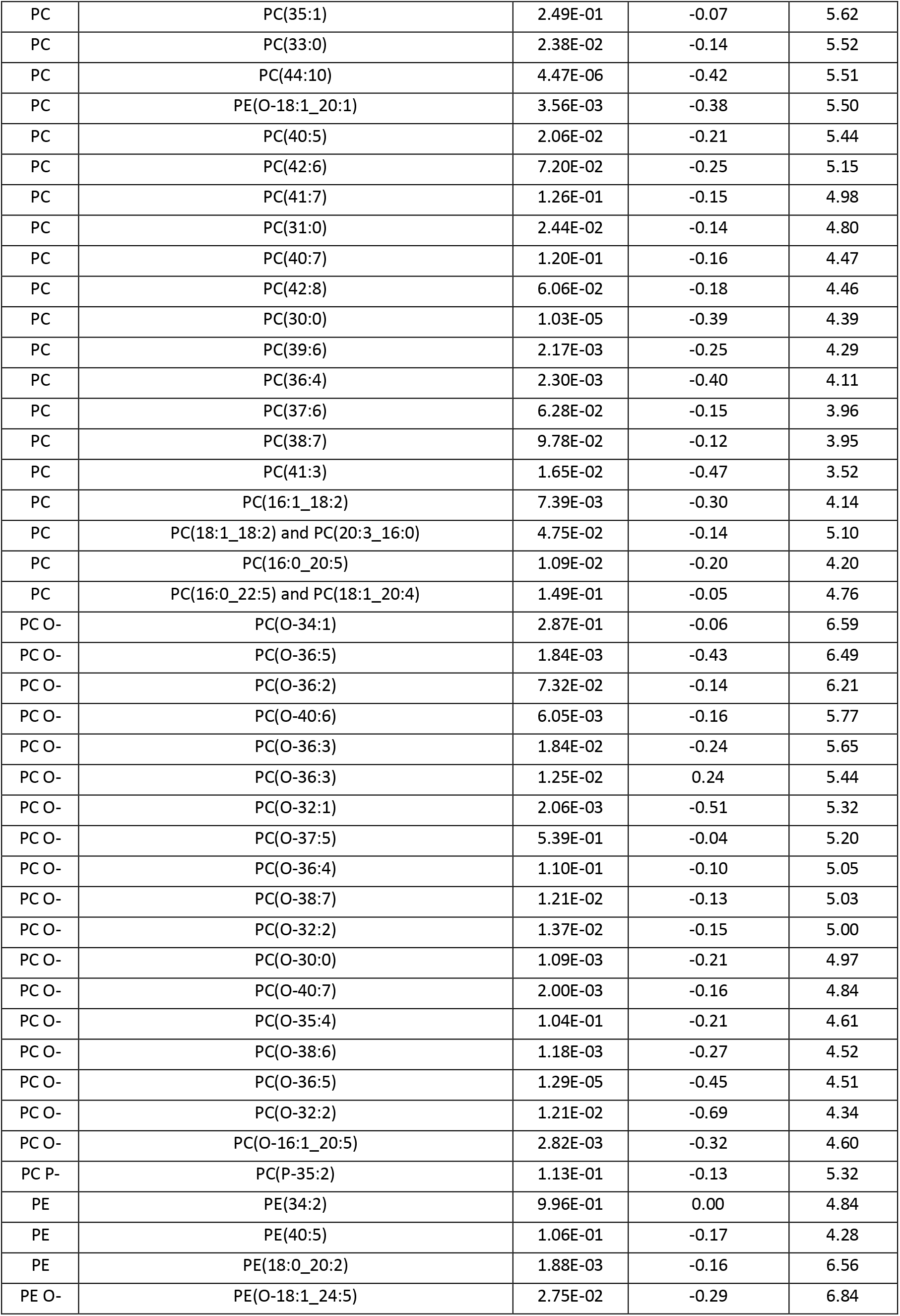
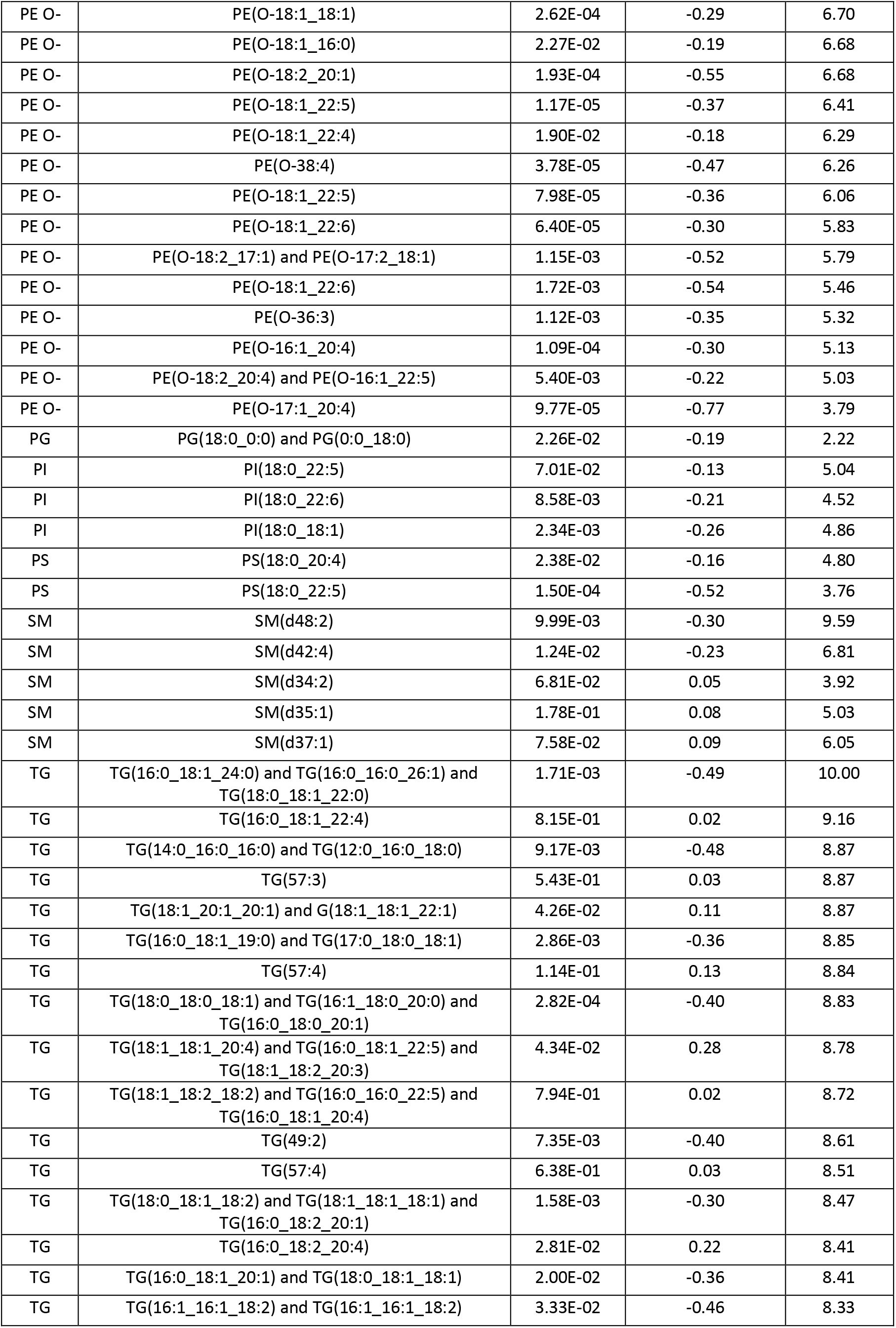
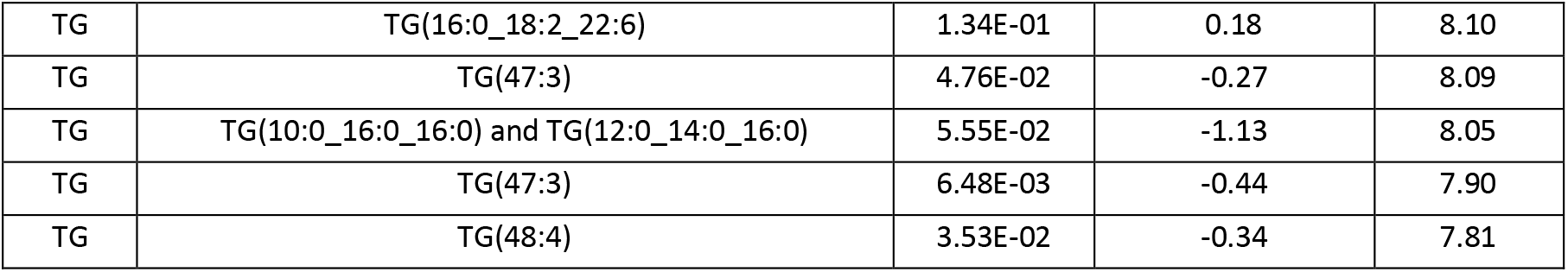
Best differential lipids for differentiating early-stage OC (I and II) *vs.* non-OC. *p*-values were calculated with Welch’s t-test. Fold changes were calculated as the base 2 logarithm of the average lipid abundance ratios for early OC *vs*. non-OC samples. A positive fold change value indicates higher levels in OC samples. Negative values indicate lower levels in OC samples. TG: Triacylglycerols, PC: Phosphatidylcholines, PC O-: Ether phosphatidylcholines, SM: Sphingomyelins, LPC: Lysophosphatidylcholines, Cer: Ceramides, PE O-: Ether phosphatidylethanolamines, Car: Carnitines, HexCer: Hexosylceramides, PE: Phosphatidylethanolamines, DG: Diacylglycerols, FA: Fatty acids, PI: Phosphatidylinositols, CE: Cholesterol esters, LPE: Lysophosphatidylethanolamines, PS: Phosphatidylserines, PG: Phosphatidylglycerols, MG: Monoradylglycerols, LPE O-: Ether Lysophosphatidylethanolamines, PS O- : Ether phosphatidylserines.

| Lipid class | Annotation | T-test p-value | Log2 fold change (OC/non-OC) | Retention Time (minutes) |
| --- | --- | --- | --- | --- |
| Car | Car(21:2) | 1.42E-03 | -0.27 | 2.61 |
| Cer | Cer(d18:1_25:1) | 5.89E-02 | 0.15 | 7.23 |
| Cer | Cer(d18:0_24:1) | 5.63E-01 | 0.07 | 7.19 |
| Cer | Cer(d18:1_25:1) | 1.30E-01 | 0.12 | 7.16 |
| Cer | Cer(d18:2_26:1) | 5.61E-03 | 0.28 | 7.12 |
| Cer | HexCer(d18:1_24:1-OH) | 2.77E-01 | 0.06 | 6.92 |
| Cer | Cer(d18:1_24:2) and Cer(d18:2_24:1) | 4.17E-03 | 0.16 | 6.83 |
| Cer | Cer(d18:1_16:0) | 8.88E-03 | 0.15 | 5.34 |
| Cer | Cer(d18:1_16:1) | 2.22E-02 | 0.17 | 5.32 |
| Cer | Cer(d18:1_25:0) | 1.47E-01 | 0.08 | 7.63 |
| Hex | Hex2Cer(d18:1_24:0) | 2.50E-01 | -0.07 | 7.00 |
| Hex | HexCer(d18:1_20:0) | 1.77E-02 | -0.15 | 6.43 |
| LPC | LPC(24:1) | 1.56E-02 | -0.21 | 3.23 |
| LPC | LPC(19:1) | 1.61E-01 | -0.10 | 2.31 |
| LPC | LPC(20:2) | 2.68E-04 | -0.24 | 2.23 |
| LPC | LPC(17:1) | 1.12E-01 | -0.20 | 1.99 |
| LPC | LPC(14:0) | 4.14E-05 | -0.45 | 1.64 |
| LPC | LPC(16:0) | 1.29E-01 | -0.06 | 2.15 |
| MG | MG(18:1_0:0_0:0) | 2.62E-01 | 0.23 | 2.55 |
| PC | PC(38:0) | 2.45E-01 | -0.18 | 7.08 |
| PC | PC(36:0) | 2.02E-02 | -0.15 | 6.76 |
| PC | PC(38:1) | 1.73E-03 | -0.22 | 6.75 |
| PC | PC(37:3) | 3.61E-05 | -0.63 | 6.66 |
| PC | PC(42:4) | 8.28E-01 | -0.01 | 6.60 |
| PC | PC(37:1) | 5.56E-03 | -0.19 | 6.59 |
| PC | PC(39:4) | 1.90E-01 | -0.10 | 6.40 |
| PC | PC(42:6) | 1.16E-03 | -0.34 | 6.16 |
| PC | PC(40:4) | 5.64E-01 | -0.05 | 6.06 |
| PC | PC(42:8) | 1.70E-02 | -0.34 | 5.75 |
| PC | PC (35:2) | 2.69E-04 | -0.42 | 5.72 |
| PC | PC(42:6) | 1.64E-02 | -0.29 | 5.70 |
| PC | PC(35:1) | 2.49E-01 | -0.07 | 5.62 |
| PC | PC(33:0) | 2.38E-02 | -0.14 | 5.52 |
| PC | PC(44:10) | 4.47E-06 | -0.42 | 5.51 |
| PC | PE(O-18:1_20:1) | 3.56E-03 | -0.38 | 5.50 |
| PC | PC(40:5) | 2.06E-02 | -0.21 | 5.44 |
| PC | PC(42:6) | 7.20E-02 | -0.25 | 5.15 |
| PC | PC(41:7) | 1.26E-01 | -0.15 | 4.98 |
| PC | PC(31:0) | 2.44E-02 | -0.14 | 4.80 |
| PC | PC(40:7) | 1.20E-01 | -0.16 | 4.47 |
| PC | PC(42:8) | 6.06E-02 | -0.18 | 4.46 |
| PC | PC(30:0) | 1.03E-05 | -0.39 | 4.39 |
| PC | PC(39:6) | 2.17E-03 | -0.25 | 4.29 |
| PC | PC(36:4) | 2.30E-03 | -0.40 | 4.11 |
| PC | PC(37:6) | 6.28E-02 | -0.15 | 3.96 |
| PC | PC(38:7) | 9.78E-02 | -0.12 | 3.95 |
| PC | PC(41:3) | 1.65E-02 | -0.47 | 3.52 |
| PC | PC(16:1_18:2) | 7.39E-03 | -0.30 | 4.14 |
| PC | PC(18:1_18:2) and PC(20:3_16:0) | 4.75E-02 | -0.14 | 5.10 |
| PC | PC(16:0_20:5) | 1.09E-02 | -0.20 | 4.20 |
| PC | PC(16:0_22:5) and PC(18:1_20:4) | 1.49E-01 | -0.05 | 4.76 |
| PC O- | PC(O-34:1) | 2.87E-01 | -0.06 | 6.59 |
| PC O- | PC(O-36:5) | 1.84E-03 | -0.43 | 6.49 |
| PC O- | PC(O-36:2) | 7.32E-02 | -0.14 | 6.21 |
| PC O- | PC(O-40:6) | 6.05E-03 | -0.16 | 5.77 |
| PC O- | PC(O-36:3) | 1.84E-02 | -0.24 | 5.65 |
| PC O- | PC(O-36:3) | 1.25E-02 | 0.24 | 5.44 |
| PC O- | PC(O-32:1) | 2.06E-03 | -0.51 | 5.32 |
| PC O- | PC(O-37:5) | 5.39E-01 | -0.04 | 5.20 |
| PC O- | PC(O-36:4) | 1.10E-01 | -0.10 | 5.05 |
| PC O- | PC(O-38:7) | 1.21E-02 | -0.13 | 5.03 |
| PC O- | PC(O-32:2) | 1.37E-02 | -0.15 | 5.00 |
| PC O- | PC(O-30:0) | 1.09E-03 | -0.21 | 4.97 |
| PC O- | PC(O-40:7) | 2.00E-03 | -0.16 | 4.84 |
| PC O- | PC(O-35:4) | 1.04E-01 | -0.21 | 4.61 |
| PC O- | PC(O-38:6) | 1.18E-03 | -0.27 | 4.52 |
| PC O- | PC(O-36:5) | 1.29E-05 | -0.45 | 4.51 |
| PC O- | PC(O-32:2) | 1.21E-02 | -0.69 | 4.34 |
| PC O- | PC(O-16:1_20:5) | 2.82E-03 | -0.32 | 4.60 |
| PC P- | PC(P-35:2) | 1.13E-01 | -0.13 | 5.32 |
| PE | PE(34:2) | 9.96E-01 | 0.00 | 4.84 |
| PE | PE(40:5) | 1.06E-01 | -0.17 | 4.28 |
| PE | PE(18:0_20:2) | 1.88E-03 | -0.16 | 6.56 |
| PE O- | PE(O-18:1_24:5) | 2.75E-02 | -0.29 | 6.84 |
| PE O- | PE(O-18:1_18:1) | 2.62E-04 | -0.29 | 6.70 |
| PE O- | PE(O-18:1_16:0) | 2.27E-02 | -0.19 | 6.68 |
| PE O- | PE(O-18:2_20:1) | 1.93E-04 | -0.55 | 6.68 |
| PE O- | PE(O-18:1_22:5) | 1.17E-05 | -0.37 | 6.41 |
| PE O- | PE(O-18:1_22:4) | 1.90E-02 | -0.18 | 6.29 |
| PE O- | PE(O-38:4) | 3.78E-05 | -0.47 | 6.26 |
| PE O- | PE(O-18:1_22:5) | 7.98E-05 | -0.36 | 6.06 |
| PE O- | PE(O-18:1_22:6) | 6.40E-05 | -0.30 | 5.83 |
| PE O- | PE(O-18:2_17:1) and PE(O-17:2_18:1) | 1.15E-03 | -0.52 | 5.79 |
| PE O- | PE(O-18:1_22:6) | 1.72E-03 | -0.54 | 5.46 |
| PE O- | PE(O-36:3) | 1.12E-03 | -0.35 | 5.32 |
| PE O- | PE(O-16:1_20:4) | 1.09E-04 | -0.30 | 5.13 |
| PE O- | PE(O-18:2_20:4) and PE(O-16:1_22:5) | 5.40E-03 | -0.22 | 5.03 |
| PE O- | PE(O-17:1_20:4) | 9.77E-05 | -0.77 | 3.79 |
| PG | PG(18:0_0:0) and PG(0:0_18:0) | 2.26E-02 | -0.19 | 2.22 |
| PI | PI(18:0_22:5) | 7.01E-02 | -0.13 | 5.04 |
| PI | PI(18:0_22:6) | 8.58E-03 | -0.21 | 4.52 |
| PI | PI(18:0_18:1) | 2.34E-03 | -0.26 | 4.86 |
| PS | PS(18:0_20:4) | 2.38E-02 | -0.16 | 4.80 |
| PS | PS(18:0_22:5) | 1.50E-04 | -0.52 | 3.76 |
| SM | SM(d48:2) | 9.99E-03 | -0.30 | 9.59 |
| SM | SM(d42:4) | 1.24E-02 | -0.23 | 6.81 |
| SM | SM(d34:2) | 6.81E-02 | 0.05 | 3.92 |
| SM | SM(d35:1) | 1.78E-01 | 0.08 | 5.03 |
| SM | SM(d37:1) | 7.58E-02 | 0.09 | 6.05 |
| TG | TG(16:0_18:1_24:0) and TG(16:0_16:0_26:1) and TG(18:0_18:1_22:0) | 1.71E-03 | -0.49 | 10.00 |
| TG | TG(16:0_18:1_22:4) | 8.15E-01 | 0.02 | 9.16 |
| TG | TG(14:0_16:0_16:0) and TG(12:0_16:0_18:0) | 9.17E-03 | -0.48 | 8.87 |
| TG | TG(57:3) | 5.43E-01 | 0.03 | 8.87 |
| TG | TG(18:1_20:1_20:1) and G(18:1_18:1_22:1) | 4.26E-02 | 0.11 | 8.87 |
| TG | TG(16:0_18:1_19:0) and TG(17:0_18:0_18:1) | 2.86E-03 | -0.36 | 8.85 |
| TG | TG(57:4) | 1.14E-01 | 0.13 | 8.84 |
| TG | TG(18:0_18:0_18:1) and TG(16:1_18:0_20:0) and TG(16:0_18:0_20:1) | 2.82E-04 | -0.40 | 8.83 |
| TG | TG(18:1_18:1_20:4) and TG(16:0_18:1_22:5) and TG(18:1_18:2_20:3) | 4.34E-02 | 0.28 | 8.78 |
| TG | TG(18:1_18:2_18:2) and TG(16:0_16:0_22:5) and TG(16:0_18:1_20:4) | 7.94E-01 | 0.02 | 8.72 |
| TG | TG(49:2) | 7.35E-03 | -0.40 | 8.61 |
| TG | TG(57:4) | 6.38E-01 | 0.03 | 8.51 |
| TG | TG(18:0_18:1_18:2) and TG(18:1_18:1_18:1) and TG(16:0_18:2_20:1) | 1.58E-03 | -0.30 | 8.47 |
| TG | TG(16:0_18:2_20:4) | 2.81E-02 | 0.22 | 8.41 |
| TG | TG(16:0_18:1_20:1) and TG(18:0_18:1_18:1) | 2.00E-02 | -0.36 | 8.41 |
| TG | TG(16:1_16:1_18:2) and TG(16:1_16:1_18:2) | 3.33E-02 | -0.46 | 8.33 |
| TG | TG(16:0_18:2_22:6) | 1.34E-01 | 0.18 | 8.10 |
| TG | TG(47:3) | 4.76E-02 | -0.27 | 8.09 |
| TG | TG(10:0_16:0_16:0) and TG(12:0_14:0_16:0) | 5.55E-02 | -1.13 | 8.05 |
| TG | TG(47:3) | 6.48E-03 | -0.44 | 7.90 |
| TG | TG(48:4) | 3.53E-02 | -0.34 | 7.81 |

**Table S8.** Significantly altered lipids derived from stage stratified analysis. *p*-values were calculated with Welch’s t-test. Fold changes were calculated as the base 2 logarithm of the average lipid abundance ratios for OC *vs*. non-OC samples. A positive fold change value indicates higher levels in OC samples. Negative values indicate lower levels in OC samples. Lipids with significant differences in the serum abundance (log2 fold change higher than 0.5 or lower than -0.5 and *p*-value less than 0.05) between (A) early-stage OC (I and II) *vs*. non-OC (B) advanced stage OC (III and IV) *vs*. non-OC. TG: Triacylglycerols, PC: Phosphatidylcholines, PC O-: Ether phosphatidylcholines, SM: Sphingomyelins, LPC: Lysophosphatidylcholines, Cer: Ceramides, PE O-: Ether phosphatidylethanolamines, PE: Phosphatidylethanolamines, PS: Phosphatidylserines.

| A. Early-Stage OC vs. Non-OC |  |  |
| --- | --- | --- |
| Annotation | T-test p-value | Log<sub>2</sub> Fold change |
| PC(37:3) | 3.61E-05 | -0.63 |
| PC(O-32:1) | 2.06E-03 | -0.51 |
| PC(O-32:2) | 1.21E-02 | -0.69 |
| PE(O-17:1_20:4) | 9.77E-05 | -0.77 |
| PE(O-18:1_22:6) | 1.72E-03 | -0.54 |
| PE(O-18:2_17:1) and PE(O-17:2_18:1) | 1.15E-03 | -0.52 |
| PE(O-18:2_20:1) | 1.93E-04 | -0.55 |
| PS(18:0_22:5) | 1.50E-04 | -0.52 |
| B. Advanced Stage OC vs. Non-OC |  |  |
| Cer(d18:0_18:1) | 6.17E-12 | 0.77 |
| Cer(d18:1_24:2) and Cer(d18:2_24:1) | 2.38E-06 | 0.53 |
| Cer(d18:2_26:1) | 7.12E-10 | 0.58 |
| LPC(14:0) | 5.14E-08 | -0.60 |
| PC (35:2) | 1.07E-09 | -0.72 |
| PC(28:0) | 3.04E-04 | -0.98 |
| PC(36:4) | 1.24E-04 | -0.61 |
| PC(40:8) | 1.02E-11 | -0.51 |
| PC(41:3) | 1.65E-02 | -0.54 |
| PC(O-18:1_22:6) and PC(O-22:7_18:0) | 4.60E-09 | -0.71 |
| PC(O-32:1) | 7.42E-06 | -0.72 |
| PC(O-36:5) | 1.82E-08 | -0.77 |
| PE(O-16:1_20:4) | 1.85E-10 | -0.53 |
| PE(O-17:1_20:4) | 5.85E-06 | -0.86 |
| PE(O-18:1_22:5) | 1.31E-07 | -0.54 |
| PE(O-18:1_22:5) | 7.08E-08 | -0.51 |
| PE(O-18:1_24:5) | 7.69E-05 | -0.51 |
| PE(O-18:2_17:1) and PE(O-17:2_18:1) | 3.87E-10 | -1.04 |
| PE(O-36:3) | 2.59E-09 | -0.64 |
| PE(O-38:4) | 8.46E-11 | -0.79 |
| PS(18:0_22:5) | 4.33E-08 | -0.69 |
| TG(14:0_16:0_16:0) and TG(12:0_16:0_18:0) | 4.32E-05 | -0.73 |

**Table S9.** Machine learning performance using the 10-lipid panel and the imbalanced dataset. Classification metrics for OC *vs*. non-OC samples in the training set and OC *vs.* non-OC in the test are shown. Five different machine learning models – random forests, logistic regression, k-Nearest Neighbor (KNN), support vector machines (SVM) and voting classifier, which is an ensemble of the four listed classifiers — were used for classification.

|  |  | **Training set**<br>OC: n= 144,<br>Non-OC: n= 83 | **Test set**<br>OC: n= 64,<br>Non-OC: n= 34 |
| --- | --- | --- | --- |
| **Random forests** | AUC | 0.72 ±0.12 | 0.75 |
|  | Accuracy | 0.71 ±0.07 | 0.74 |
|  | Sensitivity | 0.82 ±0.08 | 0.81 |
|  | Specificity | 0.52 ±0.19 | 0.62 |
| **Logistic regression** | AUC | 0.76 ±0.10 | 0.80 |
|  | Accuracy | 0.72 ±0.08 | 0.76 |
|  | Sensitivity | 0.84 ±0.11 | 0.86 |
|  | Specificity | 0.50 ±0.18 | 0.56 |
| **k-NN** | AUC | 0.67 ±0.10 | 0.70 |
|  | Accuracy | 0.70 ±0.07 | 0.68 |
|  | Sensitivity | 0.85 ±0.08 | 0.78 |
|  | Specificity | 0.46 ±0.13 | 0.50 |
| **SVM** | AUC | 0.73 ±0.12 | 0.78 |
|  | Accuracy | 0.72 ±0.08 | 0.73 |
|  | Sensitivity | 0.89 ±0.07 | 0.86 |
|  | Specificity | 0.43 ±0.19 | 0.50 |
| **Voting** | AUC | 0.74 ±0.11 | 0.78 |
|  | Accuracy | 0.71 ±0.07 | 0.69 |
|  | Sensitivity | 0.87 ±0.07 | 0.80 |
|  | Specificity | 0.44 ±0.19 | 0.50 |

**Table S10.**
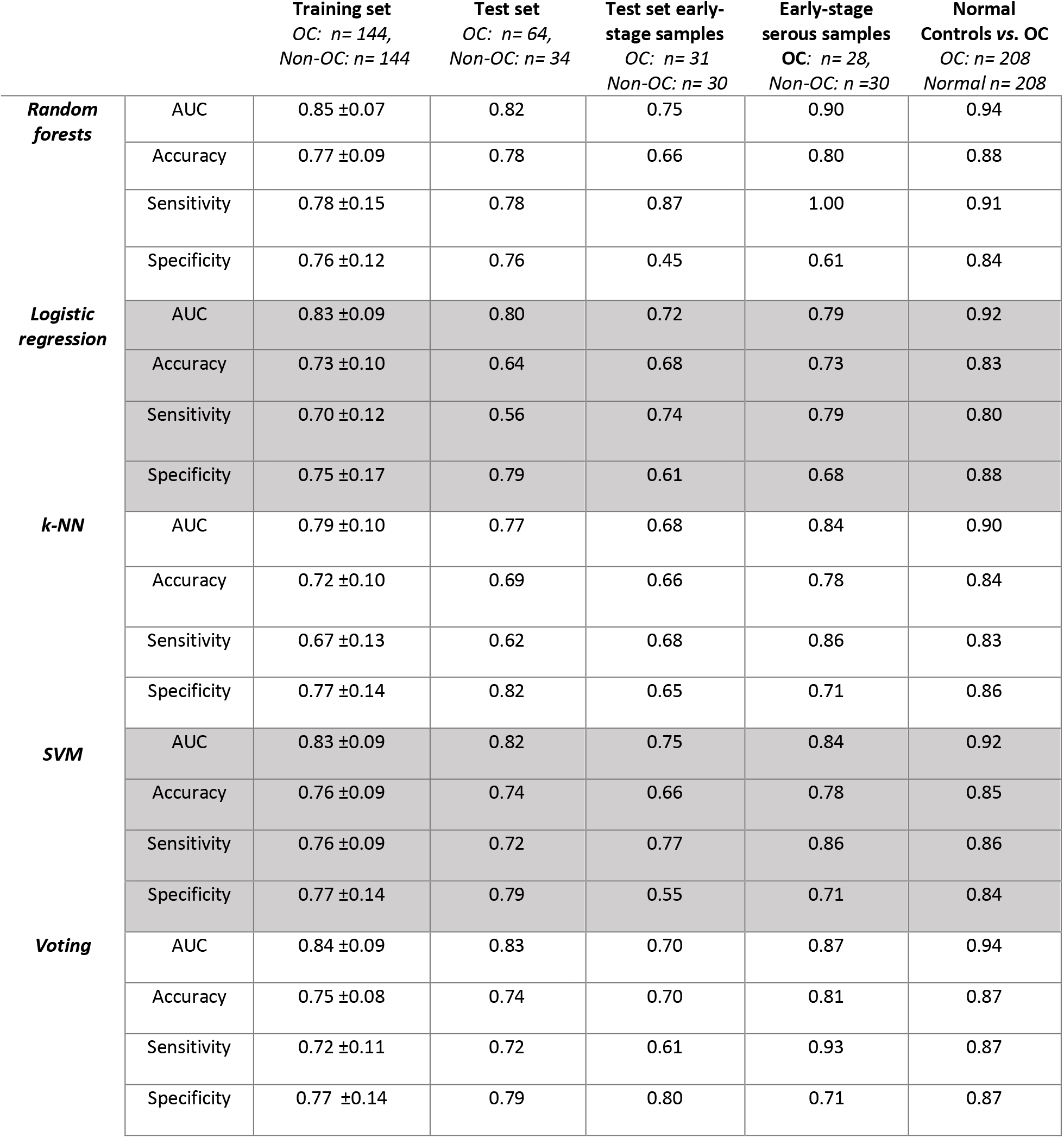
Machine learning performance for the 10-lipid panel. The training dataset was balanced with Synthetic Minority Oversampling Technique (SMOTE). Classification metrics for OC vs. non-OC samples in the training set, OC *vs.* non-OC in the test, early-stage OC (I and II) *vs.* non-OC samples from the test set, and all early-stage serous OC samples (from both training and test set) *vs*. non-OC, and normal control *vs.* OC are shown. Five different machine learning models – random forests, logistic regression, k-Nearest Neighbor (KNN), support vector machines (SVM) and voting classifier, which is an ensemble of the four listed classifiers — were used for classification.

**Table S11:**
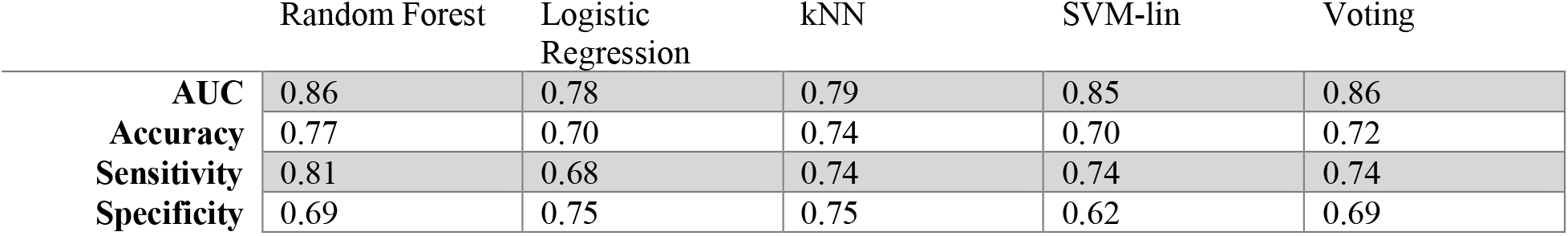
Classification performance of the 10-lipid panel for early-stage OC *vs*. benign conditions (without cervical cancer samples).

**Table S12:** 17-lipid biomarker panel including discriminant ganglioside species. Features were selected using random forest algorithm and were ranked based on the Gini index value. Proposed lipid annotation, main adduct type detected, experimental monoisotopic *m/z* value, mass error (ppm), chromatographic retention time (min), and MS annotation confidence level are shown. MS annotation level was assigned based on the following criteria: 1) MS1 and MS/MS spectrum of standard matched to the feature. 2) MS1 and MS/MS spectrum of the feature matched with library spectra 3) tentative ID assignment based on elemental formula match with literature. 4) unknowns

| Annotation | Adduct | Experimental $m/z$ | Retention Time | Mass Error (ppm) | MS Confidence Level |
| --- | --- | --- | --- | --- | --- |
| GM3(d18:0_24:2) | [M+H]<sup>+</sup> | 1263.8311 | 6.33 | 0.24 | 2 |
| GM3(d18:1_24:0) | [M+H]<sup>+</sup> | 1265.8463 | 6.70 | 0.55 | 2 |
| GM3(d34:1) | [M-H]<sup>-</sup> | 1151.7043 | 3.36 | -1.32 | 3 |
| GM1(d18:1_16:0) | [M+H]<sup>+</sup> | 1518.8530 | 3.64 | 0.26 | 2 |
| GM3(d18:1_22:1) | [M+H]<sup>+</sup> | 1235.7997 | 5.19 | 0.83 | 3 |
| GM3(d18:1_12:0) | [M+H]<sup>+</sup> | 1097.6578 | 2.80 | -0.009 | 3 |
| GM3(d18:1_16:0) | [M+H]<sup>+</sup> | 1153.7204 | 3.70 | 0.00 | 2 |
| asialo-GM1(d18:1_24:0) | [M+H]<sup>+</sup> | 1339.8835 | 6.83 | 0.76 | 2 |
| Cer(d18:1_16:0) | [M+H]<sup>+</sup> | 538.5199 | 5.33 | 0.87 | 2 |
| Cer(d18:1_25:1) | [M+H]<sup>+</sup> | 662.6451 | 7.23 | 0.77 | 2 |
| Cer(d18:2_23:0) | [M+H]<sup>+</sup> | 634.6138 | 6.93 | 0.78 | 2 |
| LPC(14:0) | [M+H]<sup>+</sup> | 468.3014 | 1.64 | 0.42 | 2 |
| PC(O-36:5) | [M+H]<sup>+</sup> | 766.5756 | 6.49 | 1.39 | 2 |
| PC(40:7) | [M+CH<sub>3</sub>COOH-H]<sup>-</sup> | 876.6123 | 5.84 | -0.14 | 2 |
| PC(O-32:2) | [M+CH<sub>3</sub>COOH-H]<sup>-</sup> | 716.5596 | 5.00 | 0.98 | 2 |
| PE(O-18:1_22:4) | [M+H]<sup>+</sup> | 758.6063 | 6.98 | -1.66 | 2 |
| PE(O-38:4) | [M+H]<sup>+</sup> | 754.5753 | 6.26 | 1.12 | 2 |
| PS(18:0_20:4) | [M+H]<sup>+</sup> | 812.5432 | 4.80 | -0.48 | 2 |

**Table S13:**
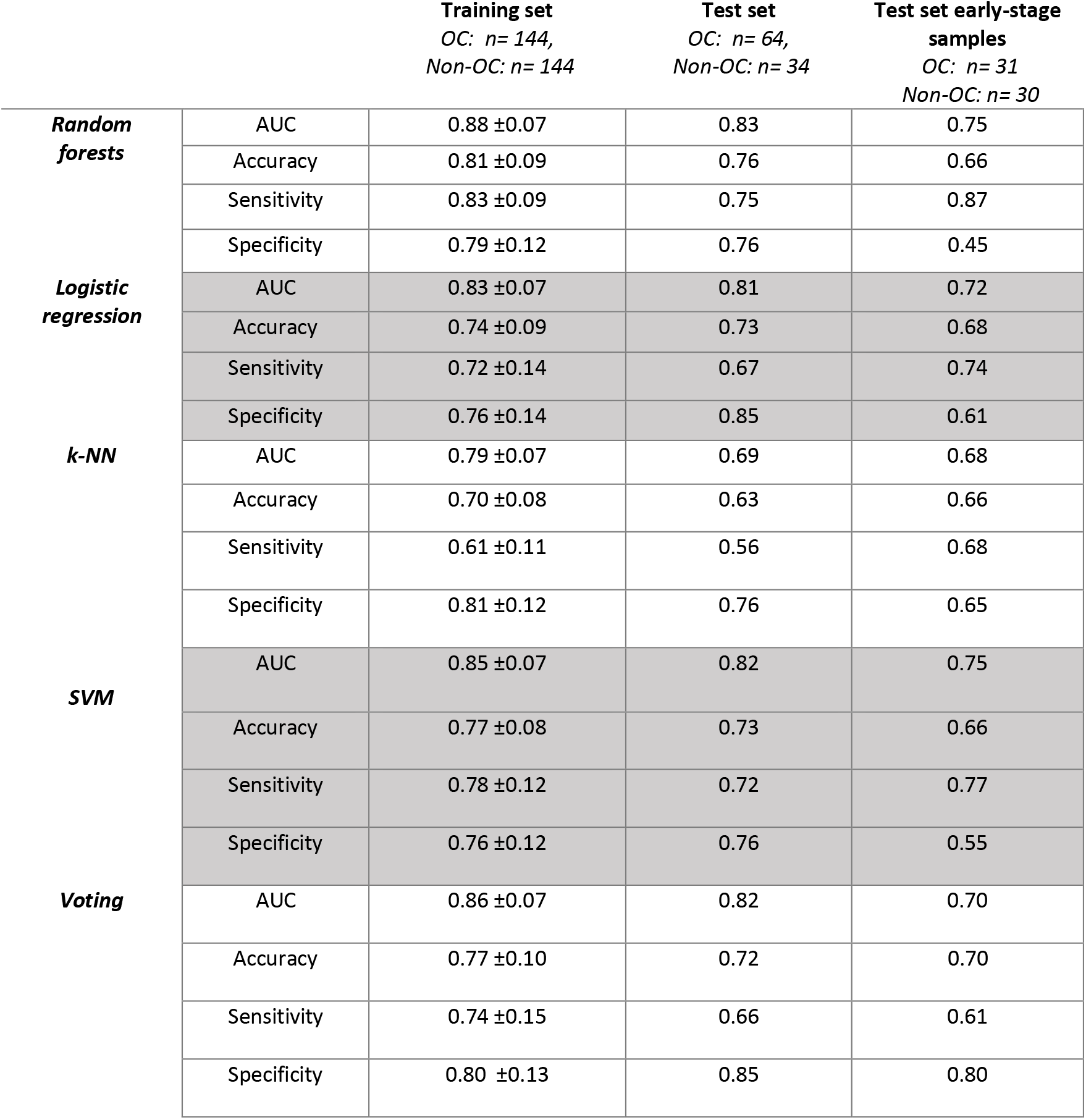
Machine learning performance for discriminant 17-lipid panel. Top 7 discriminant gangliosides were added to the 10-lipid biomarker panel to create the 17-lipid biomarker panel. The training dataset was balanced with Synthetic Minority Oversampling Technique (SMOTE). Classification metrics for OC vs non-OC samples in the training set and OC *vs.* non-OC in the test are shown. Five different machine learning models – random forests, logistic regression, k-Nearest Neighbor (KNN), support vector machines (SVM) and voting classifier, which is an ensemble of the four listed classifiers — were used for classification.

